# A tight balance of anabolic mTORC1 signaling and catabolic autophagic activity regulates zebrafish heart regeneration

**DOI:** 10.64898/2026.07.23.740244

**Authors:** Mohankrishna Dalvoy Vasudevarao, Astrid S. Pfister, Alberto Bertozzi, Thomas Kurth, Gilbert Weidinger

## Abstract

Zebrafish can regenerate the heart by proliferation of cardiomyocytes. While the innate immune response and wound re-vascularization are pre-requisites for cardiomyocyte regeneration, little is known about signals linking early injury responses with the initiations of regenerative programs in cardiomyocytes. Here we show that mTOR (mechanistic target of rapamycin) signaling is rapidly activated in response to heart injury in many cell types of the heart including endothelial cells, cardiomyocytes and macrophages, but surprisingly not in neutrophils. We find that mTORC2 regulates macrophage recruitment to the wound, while mTORC1 is required for wound debris clearance by macrophages. In addition, mTOR signaling is required for wound re-vascularization. Interestingly, it also appears to directly regulate cardiomyocyte dedifferentiation and proliferation, making mTOR signaling a central hub for regenerative responses. Anabolic mTOR signaling acts as potent inhibitor of catabolic autophagy in many systems. Yet, we observed upregulation of autophagy within border zone cardiomyocytes where mTOR signaling is active. We show that mTOR signaling limits, but does not block autophagy, and that autophagic flux is regulated by both inhibitory mTOR signaling and stimulatory JNK and MEK pathways. Our results indicate that a fine-balanced anabolic and catabolic injury response is essential for zebrafish heart regeneration. Furthermore, they reveal interesting differences in the regulation of mTOR signaling and autophagy between the regenerative zebrafish heart and non-regenerative mammalian hearts.

## Introduction

Zebrafish can efficiently regenerate heart injuries, which entails restoration of cardiomyocytes lost to injury and a lack of permanent scarring ^1–4^. Cardiomyocytes regenerate via dedifferentiation and proliferation of mature cardiomyocytes at the wound border ^5,6^. Cardiomyocyte proliferation can first be robustly detected at 3 days post injury (dpi) and peaks at 7 dpi ^1–4^. Early injury responses, in particular macrophage recruitment, re-activation of developmental gene programs in endocardium and epicardium and re-vascularization of the injured area, occur prior to and are required for subsequent cardiomyocyte proliferation ^7–10^. While many signaling pathways have been implicated in the regulation of cardiomyocyte dedifferentiation and proliferation, little is known about molecular events that link early injury responses with activation of a regenerative program in cardiomyocytes ^5,6,11–13^.

Signaling by the <u>m</u>echanistic target of rapamycin (mTOR) kinase coordinates proliferation, growth and metabolism of eukaryotic cells, but can also influence cell migration and cytoskeletal remodeling ^14^. For example, innate and adaptive immune responses require mTOR signaling ^15,16^. Classically, mTOR signaling is activated by growth factors (e.g. Insulin growth factor) in a kinase cascade involving phosphatidylinositol-4,5-bisphosphate 3-kinase (PI3K) and the serine-threonine protein kinase Akt. mTOR kinase is a component of two protein complexes, mTOR complex 1 (mTORC1) and complex 2 (mTORC2), whose activity can be disentangled by the natural product rapamycin, which only inhibits mTORC1 ^17^. mTOR signaling is required for both the adaptive and maladaptive hypertrophic response of mammalian hearts to stress ^18,19^. However, downregulation of the pathway in cardiomyocytes after acute myocardial infarction has been suggested to represent an adaptive response that promotes cardiomyocyte survival ^20,21^.

Autophagy is a cellular waste recycling process that is essential for clearance of damaged organelles and long-lived proteins and is critical for tissue maintenance and remodeling during homeostasis ^22^. Autophagy can promote cellular survival under stress, but it can also result in cell death when going unchecked or as part of a newly discovered regulated cell death program ^23–26^. Autophagy appears to be induced in mammalian cardiomyocytes after ischemic injury ^27^. However, whether it has a positive (protective) or negative effect (promoting cell death) is under debate, since evidence for both aspects has been reported ^27–34^. Autophagy has been found to play a positive, regeneration-promoting role in several other systems. For example, autophagy protects mouse muscle satellite cells from senescence, and thus is important for maintenance of the regenerative capacity of these cells ^35^. Likewise, autophagy has been reported to be required for the regeneration of zebrafish fins ^36^, of peripheral nerves ^37^, intestinal stem cells ^38^ and the liver ^39,40^. A critical feature associated with activation of mTOR signaling is the concomitant inhibition of catabolic events like autophagy, positioning mTOR signaling and autophagy at the opposing ends of the cellular metabolic spectrum ^22,41^.

In this study, we identify activation of mTOR signaling as one of the earliest known responses of zebrafish cardiomyocytes to heart injury. We show that mTOR signaling is required for many processes known to be essential for heart regeneration, including macrophage recruitment and function, wound revascularization and cardiomyocyte dedifferentiation and proliferation. Surprisingly, we found that autophagy and mTOR signaling activity can coexist in wound border cardiomyocytes, although mTOR signaling limits autophagic activity in these cells. Thus, our study indicates that an intricate balance of anabolic mTOR signaling and catabolic autophagic activities is essential for successful heart regeneration. Our results also point towards dissimilarities in the regulation and function of mTOR signaling in injured zebrafish and mammalian hearts, which might be causative for their different regenerative capacity.

## Results

### mTOR signaling is rapidly activated after heart injury in zebrafish, but not medaka cardiomyocytes

In adult mice, which fail to regenerate the heart, mTOR signaling is downregulated in cardiomyocytes following infarction ^42^. We thus wondered how pathway activity is modulated in response to injury in the regenerative adult zebrafish heart. mTOR activates p70 S6 kinase, which in turn phosphorylates ribosomal protein S6 (Rps6) at Ser 240 and 244 ^43^. We therefore used a specific antibody against Ser 240/244 of RpS6 (p-S6) as readout for mTOR signaling. Immunoblotting revealed a 4-fold upregulation of p-S6 levels in lysates of cryoinjured zebrafish ventricles (from which the wound tissue had been removed) at 3 dpi compared to uninjured hearts (**Figure 1A**). Statistical information additional to that reported in figures and figure legends can be found in **Supplementary Table 1** and in **Supplementary Table 2** for Supplementary Figures. Immunostaining of heart cryosections showed that p-S6 accumulation could already be observed at 1 dpi, where it was localized to the wound border cardiomyocytes (0-100 µm from the wound border), while we did not observe upregulation in distally located cardiomyocytes (200-300 µm from the wound border, **Figure 1B**). ∼70% of the wound border cardiomyocytes (within 100 µm of the wound border) were positive for p-S6 at 1 dpi (**Figure 1B**). At 3 dpi, p-S6 staining was particularly strong at the proximal wound border (0-50 µm, **Supplementary Figure 1A**). mTOR signaling persisted throughout the first week after injury in wound border cardiomyocytes, albeit at progressively decreasing levels, and reached baseline levels by 14 dpi (**Figure 1B**). p-S6 induction in wound border cardiomyocytes was already obvious at 6 hpi (hours post injury, **Figure 1C**), making mTOR signaling activation one of the earliest known injury responses of cardiomyocytes. Treatment of fish with rapamycin, the prototypical, highly specific inhibitor of mTORC1 ^44,45^, for 24 hours abrogated p-S6 staining in cardiomyocytes at 1 dpi, confirming that p-S6 represents a specific readout of mTOR signaling in our system (**Figure 1D**). 1 µM rapamycin was similarly effective as 100 nM when treated for 20 or 24 hours at 1 and 3 dpi, respectively (**Supplementary Figure 1B**). Thus, we continued to use 100 nM, except for experiments where our main goal was to exclude a negative result, in which case we resorted to a dose of 1 µM. While rapamycin is generally specific for mTORC1, prolonged treatment can in addition result in mTORC2 inhibition ^46^. To test the specificity of rapamycin in our system we evaluated Akt, which is phosphorylated by mTORC2 at Ser 473 ^14^. In addition to rapamycin, we utilized an ATP-site inhibitor of mTOR kinase, torin1, which inhibits both mTORC1 and mTORC2 ^47^. While torin1 significantly downregulated p-Akt levels, rapamycin treatment did not (**Figure 1E**). We conclude that rapamycin abrogated p-S6 accumulation via specific inhibition of mTORC1. Akt signaling is a prominent upstream regulator of mTORC1 in many systems ^48^. It was therefore surprising that we could not detect upregulation of p-Akt at 1 dpi, when p-S6 levels peak (**Figure 1E**, compare uninjured with DMSO groups). In fact, at 3 dpi we even observed a mild downregulation of p-Akt in untreated fish (**Figure 1F**). These data suggest that Akt does not act as predominant activator of mTOR signaling at 1 and 3 dpi in the regenerating zebrafish heart. We next asked whether mTOR signaling is also activated in other heart cell types. We could observe upregulation of p-S6 at 3 dpi in endothelial cells, labelled by the *fli1a:*eGFP^y1Tg^ transgenic line, that were located in the wound and in the wound border area (**Figure 1G**). Overall, these data show that mTOR signaling is activated early after heart injury in cardiomyocytes and endothelial cells. In contrast to zebrafish, medaka fish cannot regenerate heart injuries ^49^.

**Figure 1:**
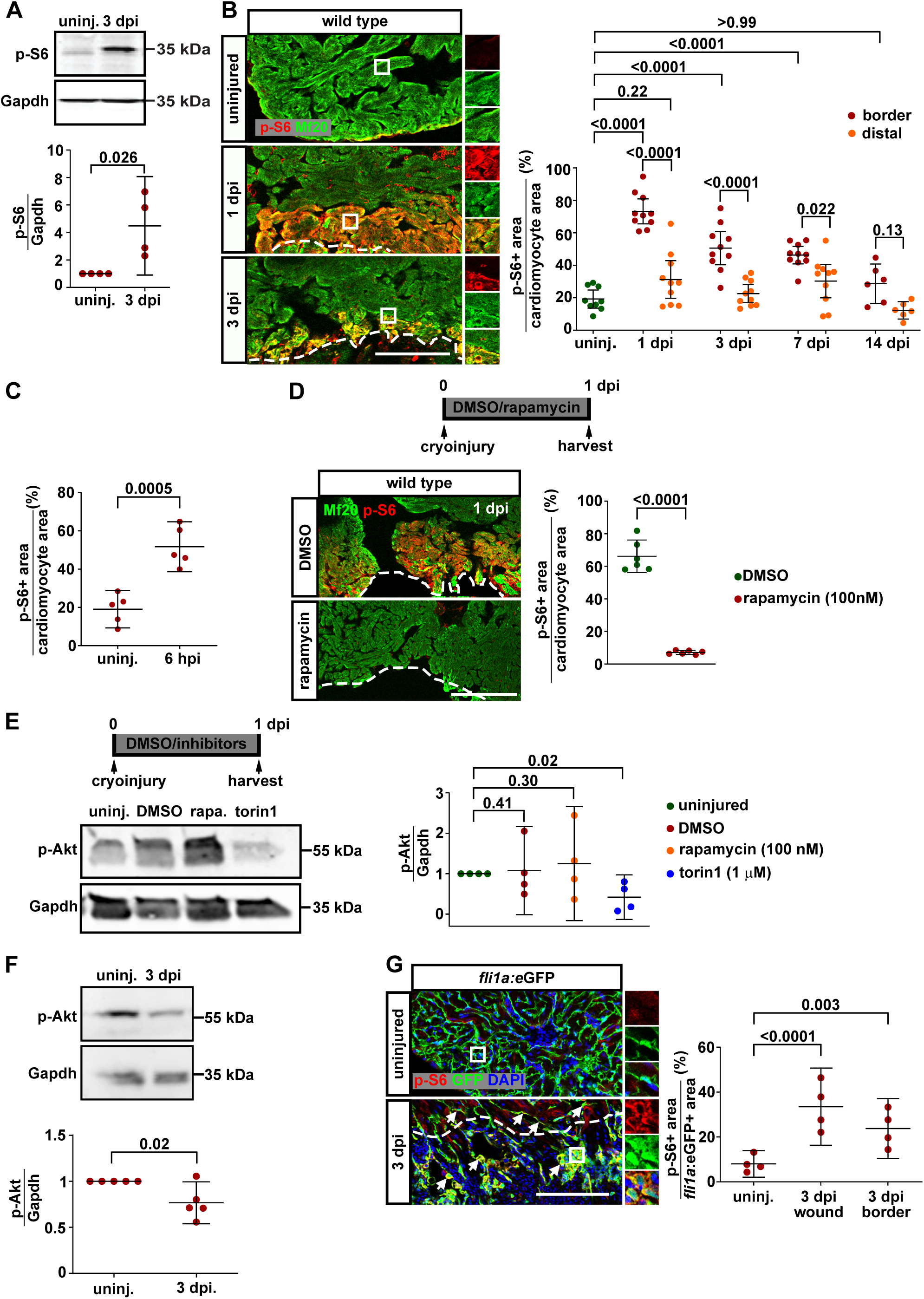
mTOR signaling is activated in cardiomyocytes and endothelial cells early after injury. (A) Immunoblotting for p-S6 reveals mTOR signaling activation in whole ventricle samples at 3 dpi relative to uninjured (uninj.) hearts. p-S6 levels relative to Gapdh and normalized to uninjured samples are plotted. Lines in graph indicate Mean ± C.I. (95%), p-value is given in the graph. For further statistical information, see **Supplementary Table 1**. (B) Immunostaining for p-S6 and Mf20 shows mTOR activation in wound border cardiomyocytes. Dashed line indicates wound border. The percentage of the Mf20^+^ area showing p-S6 staining is plotted for the wound border (0-100 μm) and distal region (200-300 μm). Scale bar, 100 µm. (C) Immunostaining for p-S6 reveals upregulation of mTOR signaling in the wound border at 6 hpi. The percentage of the Mf20^+^ area showing p-S6 staining in the wound border (0-100 µm) is plotted. (D) Treatment with 1 µM rapamycin according to the experimental scheme blocks mTOR activation in cardiomyocytes revealed by p-S6 immunostaining. Scale bar, 100 µm. (E) Immunoblotting for p-Akt comparing uninjured, 1 dpi (DMSO treated), 1 dpi (rapamycin treated, 100 nM) and 1 dpi (torin1 treated, 1 µM) ventricles indicates that mTORC1 inhibition by rapamycin is not sufficient to reduce p-Akt levels, while combined inhibition of mTORC1 and mTORC2 by torin1 is. p-Akt levels relative to Gapdh normalized to uninjured samples are plotted. (F) Immunoblotting for p-Akt comparing ventricles of uninjured and 3 dpi hearts shows downregulation of p-Akt at 3 dpi. (G) p-S6 immunostaining in *fli1a*:eGFP^y1Tg^ transgenics reveals mTOR pathway activity in endothelial cells in the wound and the wound border area at 3 dpi, which is suppressed by treatment with rapamycin for 24 hours. The GFP^+^ area showing p-S6 staining is plotted. Scale bar, 100 µm.

To test whether mTOR signaling activation is a general injury response or associated with heart regeneration, we injured medaka hearts and assayed p-S6 levels by immunostaining (**Figure 2A**). p-S6 was not upregulated in border zone cardiomyocytes (0-100 µm) in medaka hearts at 1 dpi. At 3 dpi, a few patches of cardiomyocytes became positive for p-S6 immunoreactivity, but this expression did not amount to a significant upregulation relative to uninjured hearts (**Figure 2A**). No upregulation of p-S6 was detected in cardiomyocytes located distally (200-300 µm) either at 1 or at 3 dpi (**Figure 2A**). We conclude that mTOR signaling is robustly activated in wound border cardiomyocytes in the zebrafish, but not in medaka hearts, indicating that mTOR signaling is associated with regeneration.

**Figure 2:**
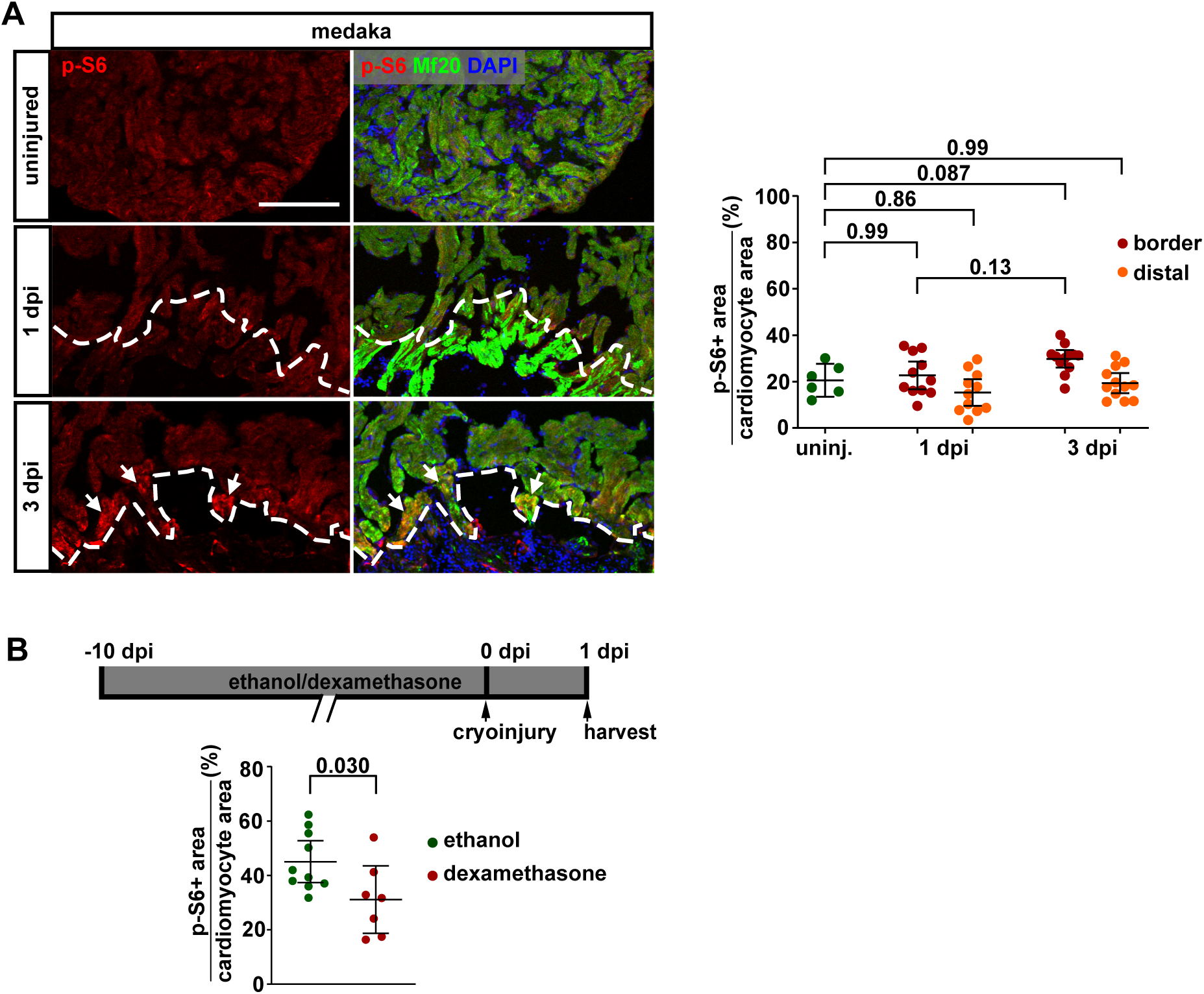
mTOR signaling is not upregulated in injured medaka hearts while inflammation is required for mTOR signaling activation in zebrafish cardiomyocytes. (A) Immunostaining for p-S6 and Mf20 shows absence of mTOR activation in wound border or distally located cardiomyocytes in medaka fish at 1 dpi and 3 dpi (uninjured vs 1 dpi border, smallest significant difference = 47%; observed difference = 10%. Note that the smallest significant difference is smaller than the biologically observed difference in Figure 1B for zebrafish, uninjured vs 1 dpi border = 281%). Scale bar, 100 µm. Lines in graph indicate Mean ± C.I. (95%). (B) Suppression of the immune response achieved by 10 days of pre-treatment with 100 µM dexamethasone reduces the area of p-S6^+^ cardiomyocytes at 1 dpi in zebrafish.

While molecular and cellular responses to heart injuries have been described to occur in the endo- and epicardium within hours, most known changes in cardiomyocytes occur later. In particular, most of the growth factor pathways that are currently known to regulate heart regeneration are not yet strongly induced at 1 dpi, but rather act during the period of maximum cardiomyocyte proliferation, which peaks at 7 dpi ^50^. We thus wondered how mTOR signaling is activated in cardiomyocytes. Inflammation occurs early after heart injury, and absence of heart regeneration is in part due to a suboptimal immune response in medaka ^9,49^. We therefore asked whether mTOR signaling activation in cardiomyocytes is dependent on the early inflammatory response. Glucocorticoids are known to inhibit heart regeneration by affecting immune cell responses ^51,52^, thus we treated fish with dexamethasone for 10 days prior to cryoinjury (**Figure 2B**). Interestingly, at 1 dpi dexamethasone treated hearts showed reduced levels of p-S6 in wound border cardiomyocytes, indicating that early activation of mTOR in the myocardium during zebrafish heart regeneration is at least partially dependent on acute inflammation and might be mediated by cytokines released by immune cells (**Figure 2B**).

### mTOR signaling is active in macrophages but not in neutrophils that are recruited to the wound

mTOR signaling has been implicated in the regulation of innate immune cell functions ^16,53,54^. Heart injury results in infiltration of neutrophils and macrophages during the first 4 dpi, which is essential for wound debris clearance and for activation of regenerative responses ^9,51,52,55^. Interestingly, we could not detect p-S6 staining in *mpx:*GFP^i114^ positive neutrophils in the wound area at 6 or 24 hpi, suggesting that neutrophils do not require mTORC1 signaling for their functions in the injured zebrafish heart (**Figure 3A**). Recruitment of *mpeg1*:eGFP^gl22^ positive macrophages to the wound could be detected by 1 dpi, which increased subsequently by 3 and 7 dpi (**Supplementary Figure 2**). Interestingly, mTOR signaling was upregulated in macrophages already by 6 hpi (**Figure 3B**). By 1 dpi ∼70% of *mpeg1*:eGFP^+^ cells were p-S6^+^, while mTOR signaling activity subsequently subsided in macrophages by 3 and 7 dpi, albeit their recruitment to the wound further increased (**Figure 3B** and **Supplementary Figure 2**).

**Figure 3:**
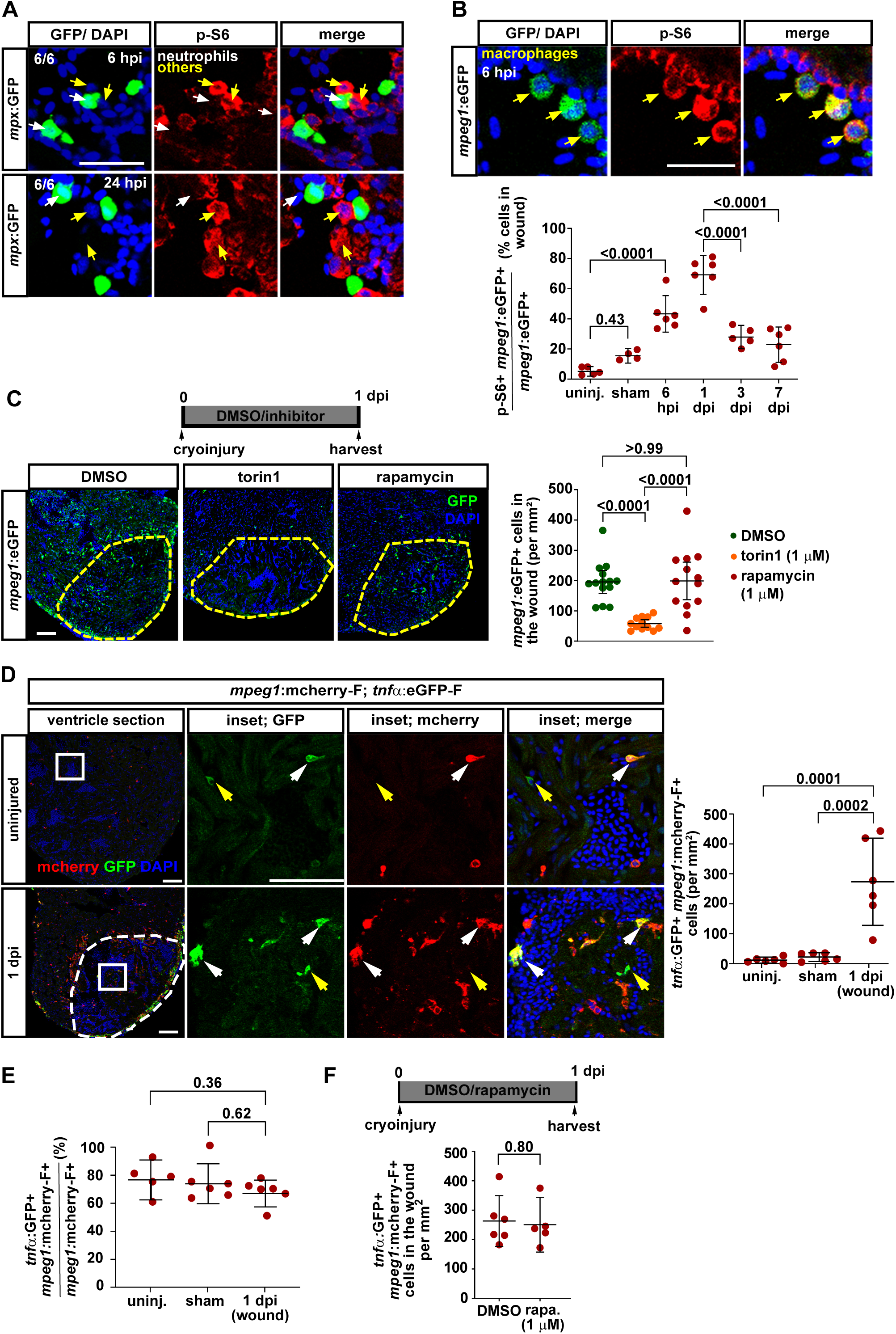
Neutrophil recruitment is independent of mTOR signaling while macrophage recruitment depends on mTORC2, but not mTORC1 activity. (A) Neutrophils (white arrows) as observed using the reporter line TgBAC(*mpx*:GFP)^i114^ do not show mTORC1 signaling activation either at 6 or 24 hours post injury in the wound area despite other cells showing p-S6 staining (yellow arrows). Scale bar, 25 µm. (B) p-S6 immunostaining in *mpeg1*:eGFP^gl22^ hearts reveals upregulation of mTOR signaling in macrophages located in the wound area, starting already at 6 hpi. By 1 dpi, nearly 70% of the GFP^+^ cells are positive for p-S6, while the number declines by 3 and 7 dpi. Scale bar, 25 µm. Lines in graph indicate Mean ± C.I. (95%). (C) Torin1 treatment (1 µM) reduces macrophage numbers recruited to the wound by 1 dpi, while rapamycin shows a much weaker effect, even at 1 µM, suggesting that mTORC2 activity regulates macrophage recruitment. Scale bar, 100 µm. (D) Tnfα^+^ pro-inflammatory macrophages, identified using *mpeg1*:mCherry-F^ump2tg^; *tnf*α:eGFP-F^ump5tg^ double transgenics, accumulate in the wound at 1 dpi. White arrows, mCherry-F^+^ GFP^+^ proinflammatory macrophages; yellow arrows, Tnfa^+^ cells which are not macrophages (GFP^+^; mCherry-F^—^ Scale bar, 100 µm (inset, 100 µm). (E) The fraction of Tnfα^+^ cells that have macrophage identity is similar in uninjured, sham and injured hearts (uninj. vs 1 dpi, smallest significant difference = 23%, observed difference = 12%; sham vs 1 dpi, smallest significant difference = 25%, observed difference = 9%). (F) Rapamycin does not affect the accumulation of pro-inflammatory macrophages (GFP^+^, mCherry- F^+^) in the wound (the smallest significant difference of 136 cells is less than the biologically observed difference of 261 cells between sham and 1 dpi in 3D).

### mTORC2 is required for macrophage recruitment, and mTORC1 for their phagocytic activity

Interestingly, torin1 strongly interfered with accumulation of macrophages in the wound at 1 dpi, while rapamycin did not, even at 1 µM, suggesting that mTORC2, but not mTORC1, is required for macrophage recruitment (**Figure 3C)**. Activated, pro-inflammatory macrophages, which can be identified by *tnfα* expression using *mpeg1*:mCherryF^ump2tg^; *tnf*α:eGFP-F^ump5tg^ transgenics ^56^, were rare in uninjured or sham injured hearts, but accumulated in the wound at 1 dpi (**Figure 3D**, white arrows). Yet, the fraction of macrophages expressing *tnf*α:eGFP-F^ump5tg^ did not differ between uninjured, sham injured or 1 dpi cryoinjured hearts, indicating that the increased number of Tnfα positive macrophages was due to recruitment of pro-inflammatory cells rather than conversion post recruitment at the wound site (**Figure 3E**). Consistent with our finding that rapamycin did not affect recruitment of the entire macrophage population, it also did not interfere with accumulation of *tnf*α:eGFP-F^ump5tg^ positive macrophages, even at 1 µM (**Figure 3F**). In summary, we conclude that mTORC2, but not mTORC1, is required for recruitment of macrophages to the wound.

Phagocytosis of tissue debris is an important function of macrophages. Cardiomyocytes in the cryoinjured area display signs of sarcomere disassembly and necrotic cell death by 1 dpi ^4^. Interestingly, while expression of GFP driven by the *myl7*:GFP^twuf1tg^ transgene was lost in these cardiomyocytes, immunoreactivity for Myosin Heavy Chain (MHC), as detected by antibodies against the MHC components Mf20 and Myl7, was still observed (**Figure 4A**). We hypothesized that the MHC positive areas in the wound represented cardiomyocyte debris, which would be cleared by macrophages. Indeed, clodronate liposome (clodrosome) injections induced severe macrophage ablation by 3 dpi (**Figure 4B**) and resulted in accumulation of Mf20 in the wound area (**Figure 4C**). Likewise, ablation of macrophages using nifurpirinol treatment of fish expressing nitroreductase under control of the *mpeg1.1* promoter (mpeg1.1:NTR-eYFPw202Tg) ^57^ (**Supplementary Figure 3**) resulted in accumulation of Mf20 in the wound at 3 dpi (**Figure 4D**). We conclude that clearance of the Mf20^+^ cardiomyocyte debris can be used as assay for macrophage phagocytic activity. While cardiomyocyte debris was cleared in DMSO treated control fish by 3 dpi, rapamycin reduced clearance at both 100 nM and 1 µM (**Figure 4E, Supplementary Figure 3B**). However, by 5 dpi, debris clearance had occurred despite continued rapamycin treatment (**Figure 4F**, red data points). Clearance was inhibited when macrophages were ablated using *mpeg1.1*:NTR-eYFP^w202Tg^ fish at 3 dpi (**Figure 4F**, yellow data points), suggesting that late phase clearance of cardiomyocyte debris is mediated by mTOR-independent macrophages. This is also supported by our observation that the number of p-S6 positive macrophages decreased from 1 to 7 dpi, while overall macrophage recruitment increased (**Figure 3B, Supplementary Figure 2**). It has been suggested that heart regeneration fails in medaka due to slower macrophage recruitment to the wound than in zebrafish ^9^. We hypothesized that this would result in reduced cardiomyocyte debris clearance in medaka fish. While the area covered by debris detected by the anti-Mf20-antibody dropped from 26% at 1 dpi to 3% at 3 dpi in zebrafish (by ∼90%), in medaka it only reduced from 41% to 20% (by ∼50% **Figure 4E and 4G**). We conclude that cardiomyocyte debris clearance is delayed in medaka fish. In summary, our data indicate that mTORC1 signaling is not required for recruitment of macrophages to the wound, but for the phagocytic activity of a subset of macrophages that is normally sufficient to clear dead cardiomyocytes by 3 dpi in zebrafish.

**Figure 4:**
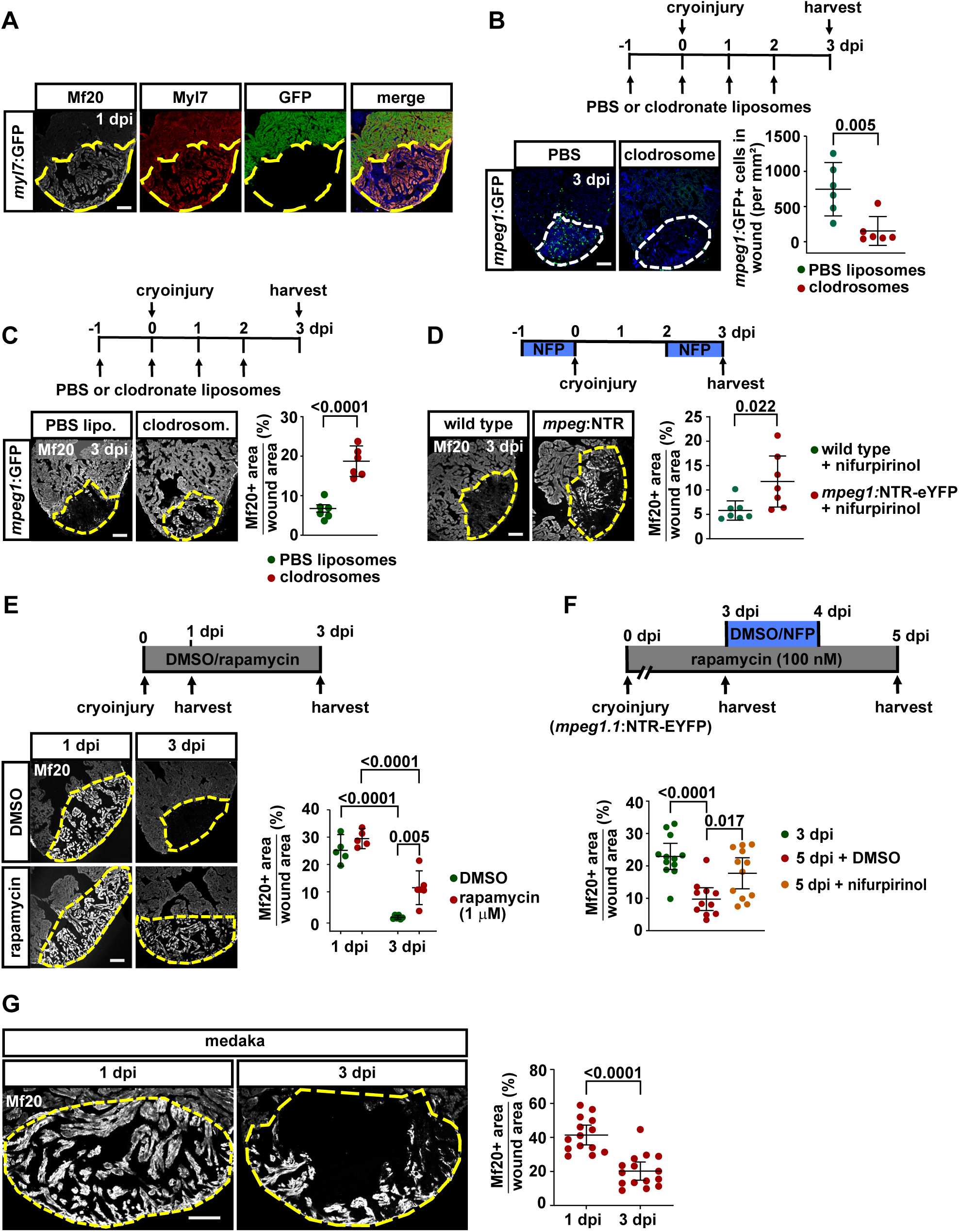
mTOR activity in macrophages is essential for cardiomyocyte debris clearance. (A) GFP fluorescence in *myl7*:GFP^twu34Tg^ transgenic hearts combined with anti MHC immunofluorescence (Mf20 and Myl7) at 1 dpi. Note Mf20 (gray) and Myl7 (red) staining in the GFP-negative wound area (dashed lines), suggesting that myocardial debris stains positive for MHC. Scale bar, 100 µm. (B) Macrophage ablation induced by daily injections of clodronate liposomes in *mpeg1.1*:eGFP^gl22^ transgenics results in drastic reduction in macrophage numbers in the wound area by 3 dpi. Scale bar, 100 µm. Lines in graph indicate Mean ± C.I. (95%). (C) Macrophage ablation results in Mf20 accumulation in the wound area at 3 dpi. Scale bar, 100 µm. (D) Macrophage ablation in *mpeg1.1*:NTR-eYFP^w202Tg^ transgenics using nifurpirinol (5 μM) results in accumulation of Mf20^+^ debris in the wound area at 3 dpi. Scale bar, 100 µm. (E) Rapamycin treatment (1 µM) causes accumulation of Mf20^+^ debris by 3 dpi. Scale bar, 100 µm. (F) *mpeg1.1*:NTR-eYFP^w202Tg^ transgenics were treated with rapamycin until 3 dpi or 5 dpi, and macrophages were ablated using nifurpirinol in one group of fish. In non-ablated hearts Mf20^+^ debris is cleared by 5 dpi (red data points), while macrophage ablation results in accumulation of debris (yellow data points). (G) Mf20 immunostaining reveals cardiomyocyte debris accumulation in injured medaka hearts at 1 and 3 dpi.

### mTOR signaling is required for wound re-vascularization, cardiomyocyte dedifferentiation and proliferation, and for scar resorption

Re-vascularization of the injured area and the wound border zone occurs quickly after heart injury and is a pre-requisite for subsequent cardiomyocyte proliferation ^8^. Since p-S6 immunoreactivity indicated that mTOR signaling is also activated after injury in endocardial cells, we asked whether it is required for wound re-vascularization. Indeed, rapamycin treatment from 0 to 3 dpi reduced proliferation of *fli1a:*eGFP^y1Tg^ labelled endothelial cells in the wound at 100 nM (**Figure 5A)** and abrogated proliferation at 1 µM (**Supplementary Figure 4A**). In addition, rapamycin impaired formation of a plexus of *fli1a:*eGFP^y1Tg^ positive cells in the injured area close to the wound border by 3 dpi (**Supplementary Figure 4B**), consistent with a previous report ^58^. We conclude that mTOR signaling is required for wound re-vascularization.

**Figure 5:**
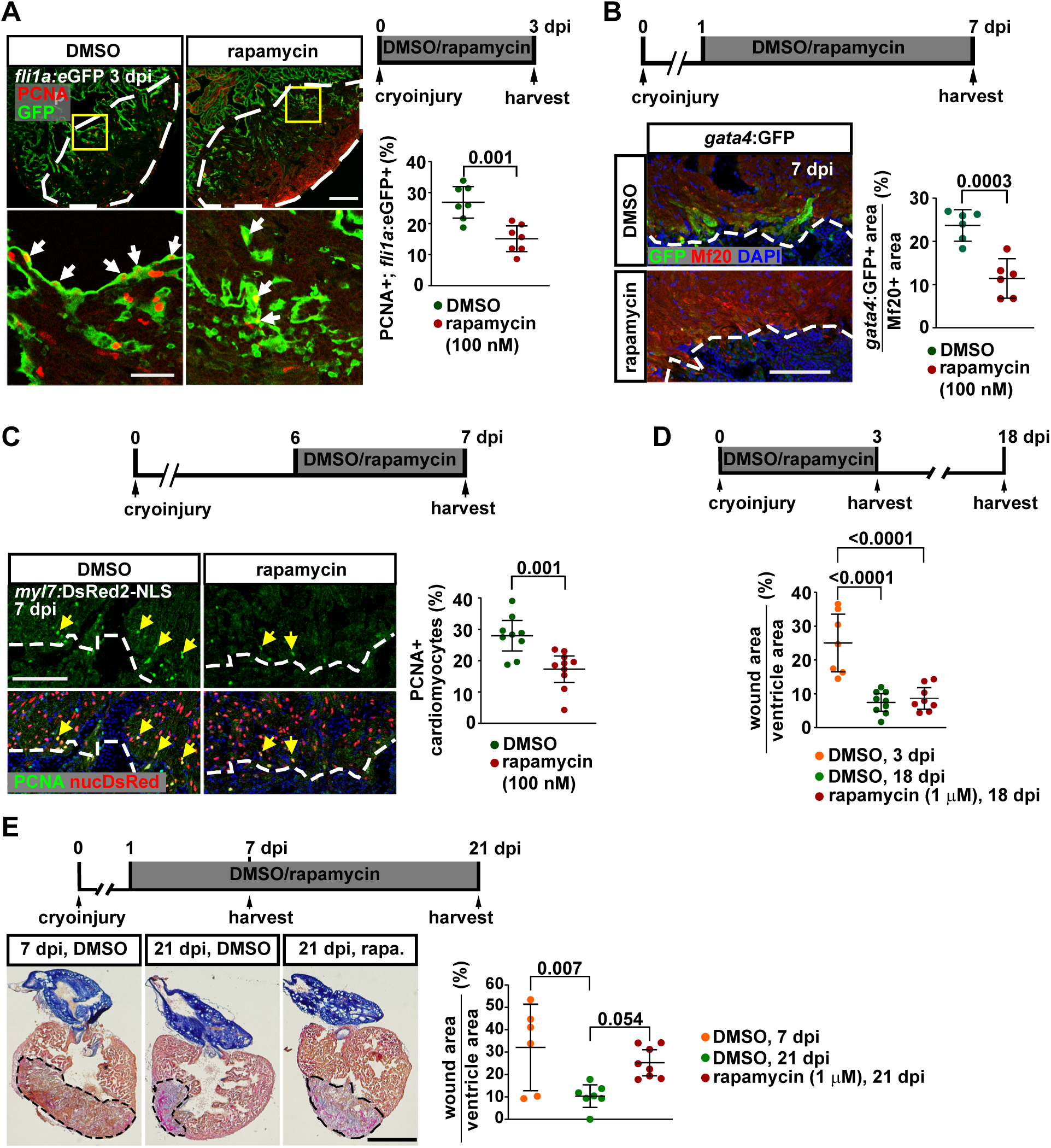
mTOR signaling is required for wound re-vascularization, cardiomyocyte dedifferentiation and proliferation, and for scar resorption. (A) Treatment with rapamycin (100 nM) reduces the number of PCNA^+^ endothelial (*flia*:eGFP^+^) cells in the wound at 3 dpi. Lines in graph indicate Mean ± C.I. (95%). Scale bar, upper panel 100 µm (lower panel 50 µm) (B) Rapamycin treatment (100 nM) downregulates injury induced expression of *gata4* regulatory sequences in Mf20^+^ cardiomyocytes of *14.8gata4*:GFP^ae1tg^ transgenic fish at 7 dpi. Size of the GFP^+^ area relative to wound border Mf20^+^ area is plotted. Scale bar, 100 µm. (C) In cryoinjured *-5.1myl7*:DsRed2-NLS^f2Tg^ transgenics rapamycin treatment (100 nM) from 6-7 dpi reduces the number of PCNA^+^ cardiomyocytes (nuclear DsRed^+^) at the wound border (0-100 µm). (D) Quantification of wound area determined by Acid fuchsin orange G (AFOG) staining in hearts that were treated from 0-3 dpi with DMSO or rapamycin (1 µM) followed by drug/solvent washout until 18 dpi. Both drug and solvent treatment resulted in wound resorption. (E) AFOG staining of heart sections reveals collagen-rich (blue) and fibrin-rich (red) wound tissue (dashed lines). Wound area has shrunken by 21 dpi in control hearts, but less so after rapamycin treatment (1 µM). Scale bar, 500 µm.

Dedifferentiation of mature cells to proliferative progenitors is increasingly being recognized as an important mechanism of regeneration in diverse systems ^59,60^. In zebrafish hearts, regenerating cardiomyocytes are derived from dedifferentiating mature cardiomyocytes ^61,62^. Cardiomyocyte dedifferentiation encompasses loss of aspects of the differentiated phenotype and gain of characteristics of an embryonic state. These are currently best characterized by loss of *myl7* promoter activity, disassembly of sarcomeres, reactivation of *gata4* regulatory sequences and re-expression of embryonic isoforms of cardiac MHC (embCMHC) detected by the N2.261 antibody ^50,62,63^. Intriguingly, rapamycin interfered with activation of *gata4* regulatory sequences observed in -*14.8gata4*:GFP*^ae1Tg^*transgenics (**Figure 5B, Supplementary Figure 4C**), reduced re-expression of embCMHC (**Supplementary Figures 4D**), reduced sarcomere disassembly in wound border cardiomyocytes (**Supplementary Figure 4E**), and decreased the downregulation of *myl7* expression as observed in *myl7*:GFP transgenic hearts (**Supplementary Figure 4F**). Thus, we conclude that mTOR signaling is required for cardiomyocyte dedifferentiation.

Dedifferentiating cardiomyocytes re-enter the cell cycle and proliferate to regenerate the myocardium ^61,62^. We thus asked whether mTOR signaling is required for cardiomyocyte proliferation. Indeed, rapamycin treatment between 0 and 3 dpi resulted in complete inhibition of cardiomyocyte cell cycle activity as observed by PCNA staining **(Supplementary Figure 5A**). Similarly, rapamycin treatment for only 24 h from 6 to 7 dpi, at the time of peak cardiomyocyte proliferation, impaired cardiomyocyte cell cycle activity, both at 100 nM (**Figure 5C)** and 1 µM (**Supplementary Figure 5B**). Importantly, this treatment did not affect endothelial plexus formation (**Supplementary Figure 5C**). We conclude that mTOR signaling is indirectly (via its early requirement for immune responses and revascularization) and directly (via its activation in cardiomyocytes) required for cardiomyocyte proliferation.

mTOR signaling is upregulated early, by 1 dpi, in cardiomyocytes and it might therefore regulate signaling events that occur at a later time point. We have previously shown that BMP signaling is upregulated in cardiomyocytes between 3 and 7 dpi and required for their dedifferentiation and proliferation ^63^. Interestingly, even at 1 µM rapamycin treatment did not result in a decrease in BMP signaling in wound border cardiomyocytes, as determined by anti-p-Smad1/5/9 (previously called Smad1/5/8) immunostaining at 3 dpi (**Supplementary Figure 5D**). Rather, a slight increase in BMP activation was observed, suggesting that these two pathways are independently regulated and that the enhanced BMP activity could be compensatory in nature.

Since we found that mTOR signaling peaks within the first 3 days after injury and has essential functions in early injury responses, we wondered whether inhibition of the pathway early after cryoinjury results in long-term defects in heart regeneration. Yet, treatment with rapamycin only from 0 to 3 dpi, followed by wash-out, did not interfere with wound resorption observed at 18 dpi, even at 1 µM (**Figure 5D**). In contrast, continuous treatment from 1 to 21 dpi resulted in significantly bigger fibrin and collagen containing wound area than in solvent-treated controls, in which the wound area had shrunk to about 1/4 of the size observed at 7 dpi (**Figure 5E**). We conclude that mTOR signaling is not only required for early injury responses, but is essential for regeneration, in particular for wound resorption, also after 3 dpi (**Figure 5E)**.

### mTOR signaling limits autophagy in border zone cardiomyocytes during heart regeneration

mTOR signaling strongly inhibits cellular catabolic processes, specifically autophagy ^14^. Considering the upregulation of mTOR signaling in wound border cardiomyocytes it could be hypothesized that autophagy is inhibited in these cells. On the other hand, injury-induced stress responses in cardiomyocytes, and in particular the process of dedifferentiation, where the sarcomeric apparatus is disassembled, might require autophagy. A recent paper reported upregulation of autophagy during zebrafish heart regeneration and suggested that this was due to downregulation of mTOR signaling, but it did not investigate whether these events occurred in the same cells ^58^. To detect autophagy, we utilized the reporter line *cmv*:eGFP-map1lc3b^zf155Tg^ (*cmv*:GFP-Lc3), which expresses GFP fused to the autophagosome marker Lc3, driven by an ubiquitously expressed *cmv* promoter. Autophagic activity results in accumulation of subcellular discrete GFP^+^ punctae ^64^. While in uninjured hearts minimal numbers of punctae per cardiomyocyte could be discerned, within 1 dpi cryoinjury resulted in accumulation of punctae in border zone cardiomyocytes (0-100 µm from the wound border), but also in cardiomyocytes in the entire basal region of the ventricle (quantified in a region 200-300 µm from the wound border, **Figure 6A**). Autophagosome numbers progressively declined during heart regeneration and reached baseline levels by 14 dpi (**Figure 6A**). Interestingly, at 1 dpi, fewer GFP-Lc3 punctae could be detected in wound border cardiomyocytes than in the distal myocardium (**Figure 6A**). While the number of punctae increased at the wound border between 1 and 3 dpi, the number at the border never reached the maximum levels observed in the distal myocardium at 1 dpi.

**Figure 6:**
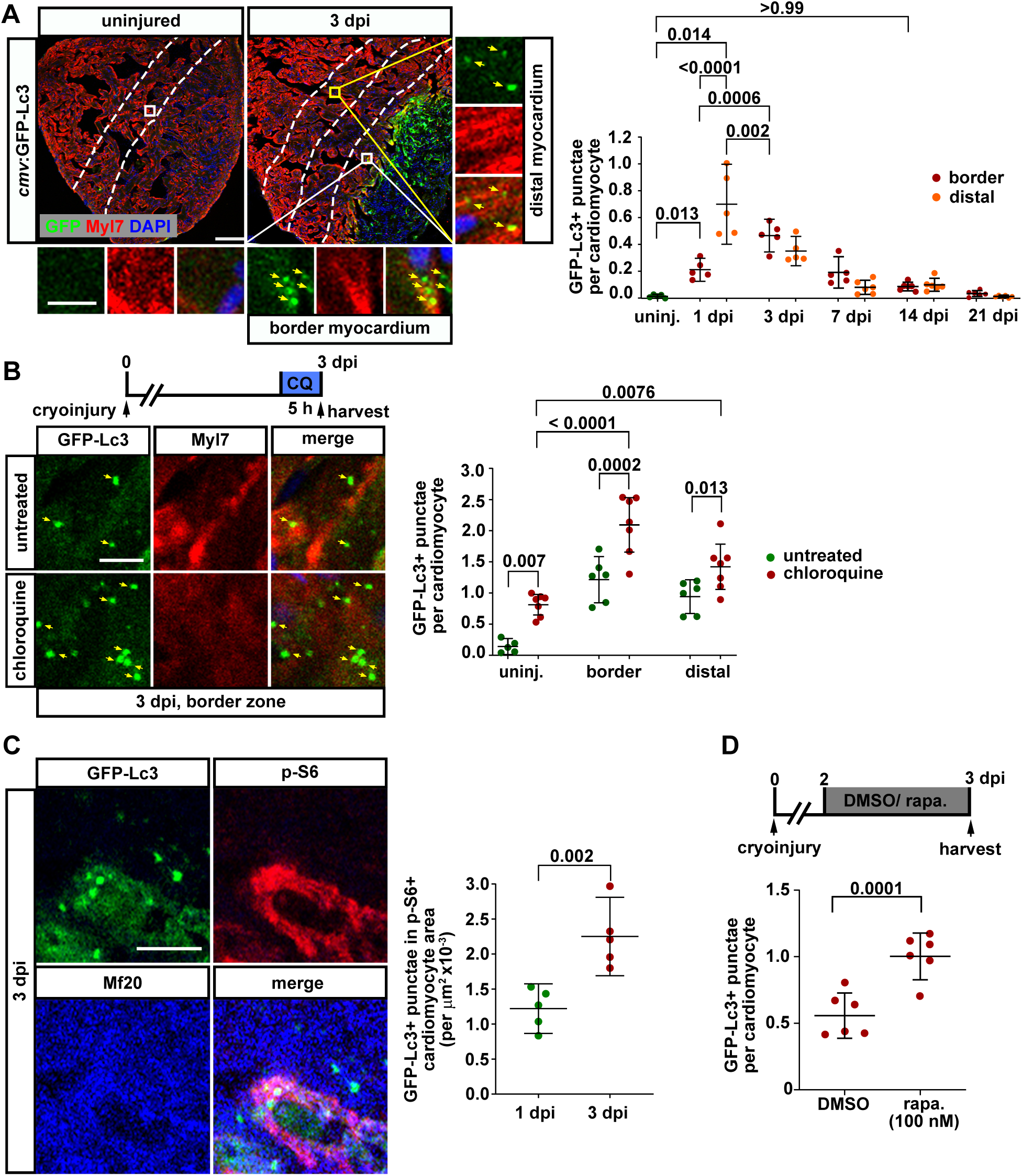
Autophagy is upregulated in cardiomyocytes during zebrafish heart regeneration, and is limited by mTOR signaling at the wound border. (A) In *cmv*:eGFP-map1lc3b^zf155Tg^ (*cmv*:GFP-Lc3) transgenics, GFP-Lc3^+^ punctae (arrows) can be detected in Myl7^+^ cardiomyocytes located at the wound border and in the distal myocardium at 3 dpi. The number of GFP^+^ punctae per cardiomyocyte in border (0-100 µm) and distal zones (200- 300 µm) is plotted. Lines in graph indicate Mean ± C.I. (95%). Scale bar, 100 µm (inset, 10 µm). (B) Treatment with 0.8 mM of the lysosomal inhibitor chloroquine (CQ) for 5 hours increases the number of GFP-Lc3^+^ punctae per Myl7^+^ cardiomyocyte at 3 dpi. Scale bar, 10 µm. (C) GFP-Lc3^+^ autophagosomes can be detected in p-S6^+^ Mf20^+^ cardiomyocytes at the wound border at 3 dpi. The number of GFP-Lc3^+^ punctae located within the p-S6^+^ Mf20^+^ double positive area within 0-100 µm of the wound border is plotted. Scale bar, 10 µm. (D) The number of GFP-Lc3^+^ punctae detected per cell in wound border (0-100 µm) Mf20^+^ cardiomyocytes at 3 dpi is increased upon treatment with 100 nM rapamycin from 2-3 dpi.

Autophagy is a dynamic process and an increase in GFP-Lc3^+^ punctae could either be due to an increase in formation or a decrease in degradation of Lc3-positive autophagic bodies ^65^. To distinguish between these possibilities, we used the lysosomal flux inhibitor chloroquine (CQ) to block degradation of autophagic bodies. An injury-induced increase of the number of GFP-Lc3^+^ punctae under such conditions would be evidence for increased formation of autophagic bodies and thus for an induction of autophagic activity. Treatment with CQ for 5 hours substantially increased the number of autophagosomes both in uninjured and injured hearts at 3 dpi, demonstrating efficient block of lysosomal function (**Figure 6B**). Interestingly, 3 dpi hearts treated with CQ showed significantly more autophagic punctae in both wound border and distal cardiomyocytes compared to uninjured fish treated with CQ, suggesting that heart injury activates formation of autophagosomes in cardiomyocytes (**Figure 6B**). Mutation of the autophagy adaptor protein p62 results in reduced levels of autophagy upon microbial infection in zebrafish ^66^. We crossed *p62*^ibl52^ fish with *cmv*:GFP-Lc3 fish to monitor autophagy in the heart. Upon cryoinjury, *p62^-/-^* fish did not show significantly different levels of autophagy compared to wild types at 3 dpi (**Supplementary Figure 6A**). However, upon treatment with CQ, *p62^-/-^* fish displayed decreased levels of GFP-Lc3 punctae compared to wild types, suggesting a decreased autophagic flux (**Supplementary Figure 6A**). These genetic data validate our assays for detection of autophagosomes and for autophagic flux inhibition in the heart. Electron microscopy revealed that autophagosomes displaying the characteristic appearance of bilamellar vesicles were not detected in uninjured hearts, but could be observed in uninjured hearts upon treatment with CQ (**Supplementary Figure 6B**). In contrast, upon cryoinjury, autophagosomes were detectable both in the border and distal myocardium in the absence of CQ treatment (**Supplementary Figure 6B**). Together, these data suggest that autophagy is quickly upregulated in cardiomyocytes in a myocardium-wide manner upon cryoinjury, while it reverts to basal levels in the second week post injury.

Due to the strong inhibitory effect of mTOR signaling on autophagy, in most systems autophagy cannot be detected if mTOR signaling is on ^67,68^. Yet, we have shown that both mTOR signaling and autophagy are activated shortly after cryoinjury in wound border cardiomyocytes. Intriguingly, we could detect co-localization of p-S6 immunoreactivity and GFP-Lc3 punctae in Mf20^+^ cardiomyocytes (**Figure 6C**). The number of these autophagosomes that appeared to escape mTOR-mediated inhibition increased between 1 and 3 dpi (**Figure 6C**). Rapamycin treatment (100 nM) for 1 day was sufficient to enhance autophagy in border zone cardiomyocytes at 3 dpi (**Figure 6D**) as did treatment with 1 µM from 4 until 7 dpi (**Supplementary Figure 6C**). Additionally, rapamycin could increase the number of GFP-Lc3^+^ punctae in the presence of CQ, confirming that mTOR signaling limits formation of autophagosomes (**Supplementary Figure 6C**). We conclude that although mTOR signaling inhibits autophagy in wound border cardiomyocytes, this inhibition is far from complete, resulting in mTOR signaling and autophagy being active in the same cells. We speculate that a fine balance between both activities is essential for allowing cardiomyocyte regeneration to occur.

### JNK and MEK signaling activate autophagy in cardiomyocytes

Stress response pathways like MEK and JNK signaling are known to induce autophagy ^69^. Indeed, zebrafish caudal fin regeneration requires autophagy, which is regulated by Erk signaling ^36^. We thus checked whether MEK and JNK signaling pathways induce autophagy during heart regeneration. Treatment with the MEK1/2 inhibitor PD 325901 or the JNK inhibitor SP 600125 resulted in downregulation of autophagy both in the wound border and the distal myocardium, as evidenced by decreased numbers of GFP-Lc3^+^ punctae (**Figure 7A, B**). Autophagic flux analysis with CQ revealed that the downregulation by both the inhibitors was due to decreased formation of autophagosomes (**Figure 7C, D**). In contrast, inhibition of MEK1/2 did not alter p-S6 activity in the border zone myocardium, indicating that mTOR signaling limits autophagy at the wound border independently of MAP kinase signaling (**Figure 7E**). Taken together our data suggest that MAP-kinase signaling and JNK signaling are required for activation of autophagy in cardiomyocytes in a myocardium-wide manner.

**Figure 7:**
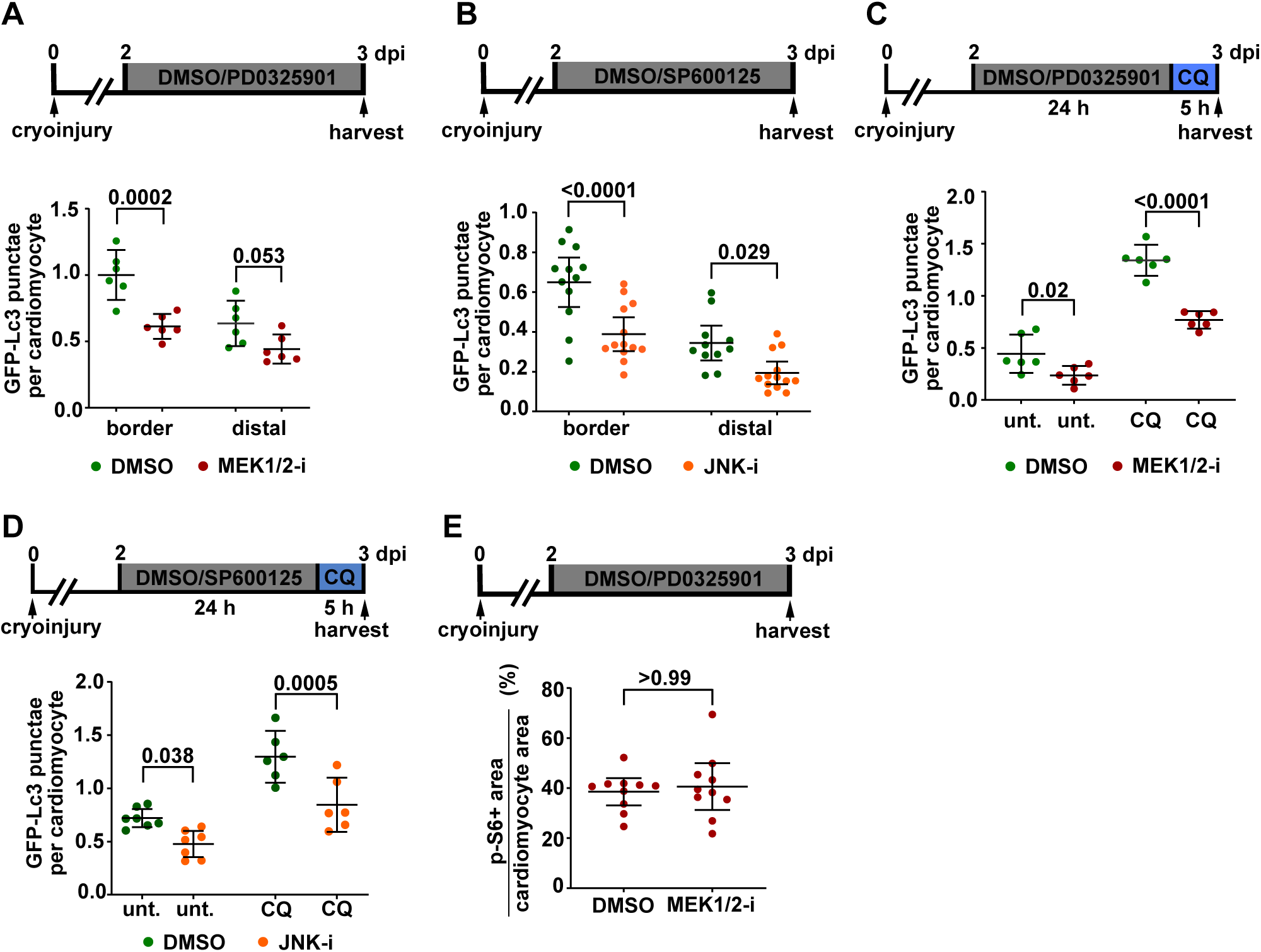
Cardiomyocyte autophagy is regulated by JNK and MEK1/2 signaling. (A) At 3 dpi, the number of GFP-Lc3^+^ punctae in Mf20^+^ cardiomyocytes located at the wound border (0-100 µm) or in distal myocardium (200-300 µm) of *cmv*:GFP-Lc3 transgenics is reduced by 24 hours of treatment with 1 µM of the MEK1/2-inhibitor PD0325901. Lines in graph indicate Mean ± C.I. (95%). (B) Treatment with 7.5 µM of the JNK inhibitor SP600125 reduces the number of GFP-Lc3^+^ autophagosomes in wound border and distal cardiomyocytes. (C) Treatment with 0.8 mM chloroquine for 5 hours in addition to treatment with PD0325901 from 2- 3 dpi shows decrease in autophagosome numbers upon chloroquine treatment in MEK1/2 inhibited hearts suggesting autophagic flux impairment. unt., untreated. (D) The JNK inhibitor SP600125 reduces autophagosome numbers upon chloroquine treatment. (E) Treatment with the MEK1/2 inhibitor PD0325901 does not change mTOR signaling in wound border (0-100 µM) cardiomyocytes as measured by the cardiomyocyte area showing p-S6 immunoreactivity at 3 dpi (smallest significant difference = 34%; observed difference = 5%. Note that the smallest significant difference is smaller than the observed difference of 38% at the wound border in Figure 7A).

## Discussion

Zebrafish hearts regenerate via dedifferentiation and proliferation of cardiomyocytes. Cardiomyocyte proliferation depends on early injury responses like immune cell infiltration and wound border re-vascularization. Here we show that mTOR signaling is a central positive regulator of zebrafish heart regeneration, since it is required for these early injury responses and in addition appears to directly regulate cardiomyocyte proliferation. Furthermore, we surprisingly found that anabolic mTOR signaling and catabolic autophagy are both active in wound border cardiomyocytes, indicating that a tight balance of these opposing processes is required for cardiomyocyte regeneration. Based on our results, we propose the following model for the role of mTOR signaling and autophagy in zebrafish heart regeneration:

mTOR signaling is upregulated by 3 dpi in endothelial / endocardial cells in the wound and the wound border and is essential for proliferation of endothelial cells and revascularization of the wound (**Figure 8A**). Surprisingly, we found no evidence for active mTOR signaling in neutrophils that are recruited to the wound (**Figure 8B**). In contrast, macrophages activate mTOR signaling, and mTORC2 is required for their recruitment to the wound, while mTORC1 regulates their phagocytic activity and is thus required for wound debris clearance (**Figure 8B**). Interestingly, mTOR signaling is also activated in wound border cardiomyocytes as early as 6 hpi, thus representing one of the earliest known responses of cardiomyocytes to heart injury. The pathway is required for cardiomyocyte dedifferentiation and proliferation (**Figure 8C**). Finally, mTOR signaling inhibition results in failure to resolve the transient scar. mTOR thus emerges as a central hub in heart regeneration.

**Figure 8:**
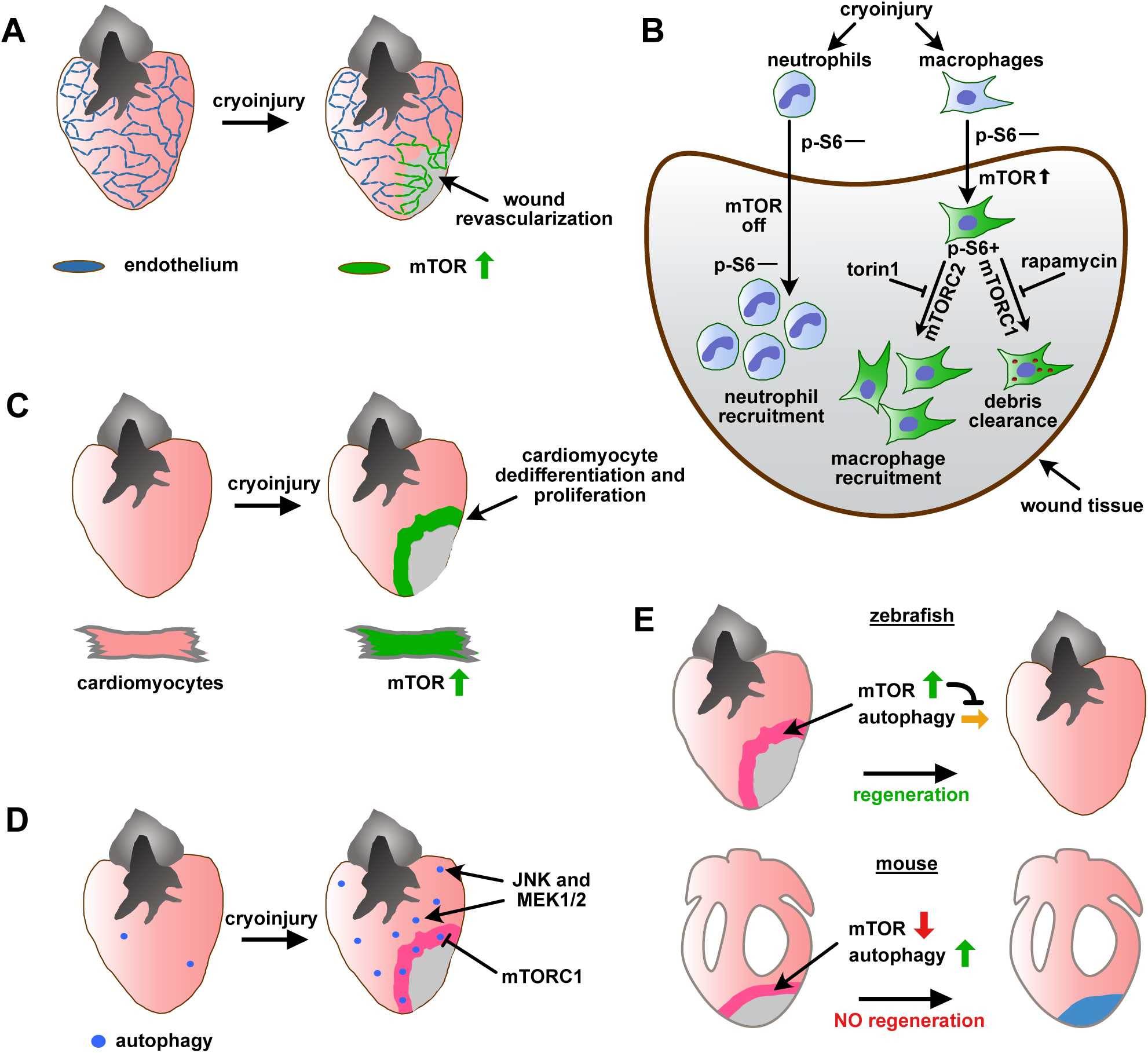
Model for mTOR-mediated regulation of a multitude of essential regenerative processes and for its interaction with cardiomyocyte autophagy. (A) mTOR signaling is upregulated after cryoinjury in endothelial cells and regulates their proliferation and the formation of a vascular plexus in the wound. (B) Neutrophils recruited to the wound are p-S6^—^ and therefore appear to function independently of mTORC1 signaling. In contrast, the majority of macrophages that accumulate in the wound are p- S6^+^ and they require mTORC2 but not mTORC1 signaling for their recruitment. In contrast, mTORC1 signaling is required for phagocytosis of wound debris by macrophages. (C) Cardiomyocytes located in the wound border upregulate mTORC1 signaling immediately after injury, which is essential for their dedifferentiation and cell cycling activity. (D) Autophagy is induced in cardiomyocytes globally in the heart within 1 day after cryoinjury, which is mediated by MEK1/2 and JNK pathways. At the wound border, a tight regulation of autophagy is maintained by mTOR signaling, which limits autophagy, and MEK1/2 and JNK pathways, which activate it. (E) Simplified model of the different roles of mTOR signaling and autophagy in adult, regenerative zebrafish and adult, non-regenerative mouse hearts. While mTOR signaling is upregulated in wound border cardiomyocytes in zebrafish and keeps autophagy in check there, mTOR signaling is downregulated in mouse, which allows for increased levels of autophagy.

Unfortunately, conditional genetic tools for cell-type specific inhibition of mTOR signaling in adult fish are not yet available. Thus, it is difficult to ascertain which of these phenotypes represent primary mTOR functions in these cell types and which are caused indirectly. Yet, the fact that mTOR signaling is active in all of the cell types for which we identified phenotypes after inhibition of the pathway argues for direct functions in macrophages, endocardial cells and cardiomyocytes. The latter is also supported by our finding that cardiomyocyte proliferation is reduced upon rapamycin treatment under conditions where endothelial cells are not affected.

Autophagy is likewise upregulated early after heart injury; in cardiomyocytes it is activated in the entire ventricle at 1 dpi **(Figure 8D**). We identified MEK1/2 and JNK signaling as pathways responsible for activation of autophagy in cardiomyocytes. Although mTOR signaling and autophagy are usually considered to be incompatible with each other, we found that cardiomyocytes at the immediate wound border, which show high levels of mTOR signaling, display autophagic activity (**Figure 8D**). Nevertheless, autophagy in these cells is limited by mTOR signaling. We thus hypothesize that the negative regulation by mTOR at the wound border assures that autophagic activity remains at low enough levels compatible with regenerative responses and cell survival. Thus, we propose that wound border cardiomyocytes require a tight balance of catabolic and anabolic pathway activities for successful heart regeneration (**Figure 8E**).

Most signaling pathways that are known to directly or indirectly regulate cardiomyocyte proliferation (Pdgf, Igf, Bmp, Tgf-β, Hh and Neuregulin) show highest activity relatively late after injury at either 3 and/or 7 dpi, at the time when cardiomyocyte proliferation peaks ^11^. Retinoic acid production is activated early after injury in the endocardium and epicardium and is required for cardiomyocyte proliferation, but it is unknown whether cardiomyocytes directly respond to it ^7,70^. Thus, with the exception of JAK-STAT signaling, which is activated in cardiomyocytes as early as 1 dpi, very little is known about cardiomyocyte responses early during regeneration ^71^. Our study identifies mTOR signaling as a pathway that is upregulated very early after injury in cardiomyocytes. mTOR signaling is best known for its role in promotion of cell growth and proliferation and it features prominently as a downstream transducer of growth factor signaling. However, at 1 dpi mTOR signaling in cardiomyocytes is unlikely downstream of growth factor signaling, since these pathways are activated at later stages. Instead, our results suggest that signals produced by inflammatory cells (e.g. cytokines) are responsible for early activation of mTOR signaling in cardiomyocytes.

mTOR signaling has important function in cells of the innate immune system ^15^ ^16^. Activation and migration of neutrophils, which are the first line of defense to tissue damage, can occur both in a mTORC1 and mTORC2 signaling dependent manner with rapamycin being a strong inhibitor of neutrophil chemotaxis ^54,72^. In addition to influencing the metabolism of immune cells, the quick response from neutrophils is partially due to mTOR dependent translation of pre-existing mRNAs like cyclooxygenase-2 (COX2), which leads to expression of prostaglandins that promote inflammation and vasodilation to attract further immune cells ^53^. Interestingly, we found that mTORC1 is not activated in neutrophils that are recruited to the wound in the injured zebrafish heart. It will be interesting to study whether zebrafish neutrophils do generally not require mTOR signaling, whether the situation in the injured heart is special, or whether the above-mentioned studies on the role of mTOR signaling in neutrophils, which were mainly performed *ex vivo*, do not reflect the situation also in other *in vivo* wound models.

mTOR signaling is a strong antagonist of autophagy ^67^ . While mTOR signaling is usually activated under growth favorable conditions by nutrients and growth factors, autophagy is a catabolic process involved in the breakdown of long-lived proteins, organelles and the recycling of metabolites under conditions of starvation and stress. Our results suggest that in wound border cardiomyocytes mTOR activity does limit autophagic activity, as it does in many other systems. Yet surprisingly, the inhibition is far from complete and many wound border cardiomyocytes display robust activation of both mTOR and autophagy. A similar observation was made in the zebrafish mutant *titaniai^s4^*^50^, which is caused by mutation in the gene periodic tryptophan protein 2 homologue (*pwp2h*), which encodes a scaffold component of the 90S pre-ribosomal particle ^73^. This mutation results in impaired ribosomal biogenesis and the mutants display a strong upregulation of both mTOR activity and autophagy. In this case, activated mTOR fails to abrogate autophagy suggesting that regulation of autophagy is complex and likely involves mTOR independent factors. Reminiscent of this is our observation that elevated mTOR activity fails to completely abrogate autophagy in the border cardiomyocytes, rather just limits it. To our knowledge, co-existence of mTOR signaling and autophagy has rarely been described in other systems and cell types, and it will be interesting to study the underlying molecular mechanisms.

A recently published study reported that autophagy is upregulated during zebrafish heart regeneration and it has used long-term rapamycin treatment as a tool to activate autophagy ^58^. While we likewise find that autophagy is activated during heart regeneration, our data is in disagreement with the study by Lavandero and colleagues ^58^ on several crucial issues. First, the authors conclude that mTOR signaling is downregulated at 3 dpi, which presumably leads to upregulation of autophagy. However, we find that mTOR signaling is strongly upregulated in several cell types at the border zone, but not in areas distal to the wound and not in some cell types like neutrophils. Chavez et al. have used Western blots of whole heart tissue for the mTOR target p4E-BP1 ^58^, while we used p-S6, for which we find upregulation both by Western blots and by immunostaining (which adds spatial resolution). Chavez et al. found that long term rapamycin treatment resulted in increased amounts of p4E-BP1, which is consistent with previous findings that p4E-BP1 can recover from prolonged rapamycin treatments ^74^. On the other hand, p-S6 is a sensitive read out for rapamycin-mediated mTORC1 suppression, irrespective of the duration of treatment ^74^. Surprisingly, Chavez et al. also report that mTORC1 inhibition by rapamycin is sufficient to decrease macrophage recruitment ^58^, while we found an effect on macrophage recruitment only when we inhibited mTORC2. While we have tried to avoid indirect and off-target effects by limiting our treatments to 24 hours, and also read out macrophage recruitment at the earliest possible time points (1 day post injury), it is possible that the long-term treatment regime used by Chavez et al. (starting at 2 days prior to injury and lasting until 5 dpi) results in indirect effects on macrophages. Finally, Chavez et al. reported that rapamycin treatment results in a minor increase in cardiomyocyte cell cycle activity ^58^, while we show a severe inhibition of cardiomyocyte proliferation. Several differences in experimental regimes could account for this discrepancy: Chavez et al. used long-term rapamycin treatment and assayed cell cycle activity at 14 dpi, while we only treated for 24 hours and analyzed hearts at the peak of cardiomyocyte cell cycle activity at 7 dpi. We assume that short-term treatments avoid complications due to indirect effects, and in fact we found that this regime did not affect wound re-vascularization, supporting a direct requirement of mTOR signaling for cardiomyocyte proliferation. Alternatively, an increase of cardiomyocyte proliferation at 14 dpi in response to long-term rapamycin treatment could be a compensatory response to a reduction of cell cycle activity at earlier time points.

The role of autophagy in the ischemic mammalian heart is far from clear ^75^. While on one hand autophagy appears to be beneficial under conditions of myocardial ischemia, enhanced autophagy is detrimental under conditions of ischemia followed by reperfusion ^28–32,34^. Interestingly, the spatiotemporal activity profile of autophagy differs between injured mouse and zebrafish hearts. In the mouse, intense but transient upregulation of autophagy is observed in the wound border 24 hours after infarction, while autophagy in remote areas increases between 1 and 3 weeks post myocardial infarction ^31^. In contrast, in zebrafish upregulation in the wound border is less than in the distal area at 1 dpi. Subsequently, in the zebrafish heart autophagy progressively decreases in the cardiomyocytes of the distal myocardium and reaches baseline values by 2 weeks post injury. Likely, the different spatiotemporal activity profile of autophagic activity reflects differential requirements for energy and organelle recycling between mouse and zebrafish cardiomyocytes, which could be related to their differential regenerative capacity.

In mammals, mTOR signaling has been shown to be essential for both development and maintenance of the heart ^20^. mTOR activity is also required for the development of adaptive hypertrophy in response to pressure overload. Interestingly, after acute myocardial infarction, mTOR signaling is downregulated in mouse cardiomyocytes and experimental activation of the pathway promotes cardiomyocyte death ^42^. Thus, it was suggested that downregulation of mTOR signaling represents an adaptive response of cardiomyocytes to myocardial infarction, partly because it results in increased autophagy, which would promote cardiomyocyte survival ^20,42^ (**Figure 8E)**. Furthermore, a downregulation of mTOR signaling has been described in several other mammalian tissues immediately post injury ^76^. This downregulation is associated with enhanced autophagy and was suggested to provide energy and raw materials for subsequent growth, where mTOR activity recurs. In contrast, we found that mTOR signaling is strongly activated early after heart injury in zebrafish, particularly in cardiomyocytes. Furthermore, our results show that activation of mTOR signaling is essential for cardiomyocyte proliferation and that prolonged rapamycin treatment is detrimental to zebrafish heart regeneration. Interestingly, in mice long-term rapamycin treatment after myocardial infarction was shown to be beneficial in terms of cardiac remodeling and cardiac function, which in turn was a result of reduced cardiac inflammation ^77^, Kanamori, 2011 #238.

Thus, our study brings to light stark differences between mouse and zebrafish with respect to the kinetics and function of mTOR signaling and the activity profile of autophagy in injured hearts (**Figure 8E**). Possibly, zebrafish cardiomyocytes at the wound border are able to proliferate and to regenerate, while mammalian cardiomyocytes in adults cannot, because zebrafish cells acquire the right balance between anabolic mTOR signaling and catabolic autophagic activity. Indeed, similar to zebrafish heart regeneration and contrary to adult mouse injury responses, neonatal mouse pups activate mTOR signaling in the wound border during regeneration ^78^. Rapamycin is a strong inhibitor of mouse neonatal cardiomyocyte proliferation as well ^78^, suggesting that activation of mTOR signaling is conserved under contexts where regeneration is the outcome. Thus, identifying the reasons underlying the differential regulation of these conserved pathways will be important for deciphering why the adult zebrafish heart can regenerate, while adult mammalian hearts cannot.

## Materials and methods

### Transgenic and mutant fish lines, cryoinjury, heat shocks, and drug treatment

All experiments using adult zebrafish have been approved by the state of Baden-Württemberg (Tierversuch Nr. 1193 and 1352) and the animal protection representative of Ulm University. The following fish lines were used: *-5.1myl7*:DsRed2-NLS^f2Tg^ ^79^, *myl7*:GFP^twu34Tg80^, *fli1a*:eGFP^y1Tg^^81^, *- 14.8gata4*:GFP^ae1tg81^, *hsp70l*:axin1^w35Tg83^, *hsp70l*:cyp26a^kn1Tg84^, *hsp70l*:nog3^fr14Tg85^, *cmv*:eGFP- map1lc3b^zf155Tg64^, TgBAC(*mpx*:GFP)^i114^ ^86^, *mpeg1*:EGFP^gl2287^, *mpeg1.1*:NTR-eYFP^w202Tg88^ and *p62*^ibl52^ mutant ^66^. Heart cryoinjuries were performed using a liquid nitrogen cooled copper filament as described previously ^63^. For sham injuries, the protocol was exactly the same as for cryoinjury except that the intervention was performed with the filament at room temperature. Heat shock overexpression lines received one heat shock at 37^°^ C for 1 hour. Fish were exposed to the following compounds in fish system water: rapamycin (Selleckchem #S1039, 1 μM or 100 nM, see figure legends), Dexamethasone (Sigma #D1756, 100 mM), Nifurpirinol (Dr. Ehrenstorfer Gmbh #DRE-C15523050, 5 μM), MEK1/2 inhibitor PD 0325901 (Selleckchem # S1036, 1 mM) and JNK inhibitor SP600125 (Selleckchem #S1460, 7.5 μM). For macrophage ablation, 10 μl of PBS containing 5 mg/ml of liposomes loaded with PBS or clodronate (Liposoma PBS-02) were injected intraperitoneally into adult zebrafish.

### Immunofluorescence and histology

For immunofluorescence and histological staining, hearts were extracted, fixed in 4% paraformaldehyde (PFA) (in phosphate buffer) at room temperature for 1 h, cryo-protected in 30% sucrose (in phosphate buffer) overnight and cryosectioned into 10 µm sections. The general protocol for immunofluorescence has been described before ^4^. Heart sections were equally distributed between 6-8 serial slides so that each slide contained sections representing all areas of the ventricle. Primary antibodies used for immunofluorescence were anti phospho-serine 240/244-ribosomal protein S6 (Cell Signaling #2215; 1:300), anti-PCNA (Dako #M0879; 1:2000), anti-GFP (Abcam #ab13970, 1:500), anti-α-actinin (Sigma #A7811; 1:500), anti-N2.261 (Developmental Studies Hybridoma Bank #AB_531790, 1:50), which detects an embryonic cardiac MHC isoform (embCMHC) in zebrafish ^50^, anti-Mf20 (which detects MHC, Developmental Studies Hybridoma Bank #AB_2147781, 1:100), anti- Myl7 (Genetex #GTX128346, 1:500), anti-Mef2c (Santa Cruz #SC313, 1:50), and anti-p-SMAD1/5/9 (Cell Signaling #13820, 1:500). Secondary antibodies conjugated to Alexa 488, 555, or 633 (Invitrogen) were used at a dilution of 1:1000. Nuclei were stained by DAPI (4’,6-diamidino-2-phenylindole). AFOG staining was performed as described before ^1^.

### Image analysis

All images of fluorescent proteins represent immunofluorescence stainings using anti-GFP antibody except for Supplementary Figures 2 and 3C, which show endogenous YFP and GFP fluorescence respectively. All images of immunofluorescence staining were acquired at 20X magnification (unless specified otherwise in the figure legends) in a single optical plane with a Leica SP5 confocal microscope. For quantifications, in injured hearts 2-3 sections displaying significant sizes of both wound and myocardium were analyzed per heart. When using transgenic reporter lines for identifying cardiomyocytes (*-5.1myl7*:DsRed2-NLS^f2Tg^ or *myl7*:GFP^twu34Tg^), upon injury, wounds were identified by the lack of reporter activity. Otherwise, wounds were identified by a combination of lack of Myosin Heavy Chain immunostaining (Mf20 or Myl7) and a qualitative change in the DAPI positive nucleus density between the wound and myocardial border. For uninjured hearts, 1-2 sections were quantified. Autophagosome counting and p-S6 quantification were performed both at the wound border (0-100 μm) and distal areas (200-300 μm). Quantifications of PCNA and nuclear DsRed were performed in cardiomyocytes situated within 100 μm from the wound border. For dedifferentiation assays, ImageJ was used for selecting the area positive for a marker or the overlap of two markers using threshold functions, and for quantifying the area, namely myocardium (Mf20, Myl7 or α-actinin staining), *14.8gata4*:GFP^ae1tg^ upregulation and upregulation of embCMHC. For *myl7*:GFP^twu34Tg^ downregulation, the GFP^low^ area was defined as the area occupied by pixels with intensity values of 11-80, while GFP^high^ was defined as pixel values 81-255. For AFOG staining, bright field imaging was performed using an Olympus microscope and quantification of wound size was performed manually using ImageJ. Serial sections of hearts were distributed among 6 slides so that every 6^th^ section was on one slide allowing for sampling 1/6^th^ of the heart for wound size. Similarly, endothelial regeneration was measured by thresholding GFP^+^ pixels in *fli1a*:eGFP^y1Tg^ hearts and measuring the area occupied by such pixels from all heart sections on the slide showing a wound.

### Statistical analyses

Lines in graphs indicate average and confidence interval (95%). Statistical significance of Western blotting datasets was calculated with the paired student’s t-test (one-tailed) and a normal distribution was assumed.. For all other tests between two groups two tailed, unpaired, student’s t-test was used. For testing significance in case of multiple groups one way ANOVA was performed which was then followed by a post hoc multiple comparison test as specified in **Supplementary Table 1** and **Supplementary Table 2**. Sample sizes are also available in Supplementary Table 1 and Supplementary Table 2. To support statements about the lack of differences between experimental groups (where p-values are > 0.05) we computed the effect size using GPower (University of Düsseldorf) with these parameters: t-test tails = two, α = 0.05, power (1-β) = 0.8 and the sample size of both groups. Then, the smallest difference that would have been significant was calculated based on the effect size and the Means, SDs and sample sizes of the experimental groups. These smallest significant differences and the observed differences are reported in the Figure legends. If we considered the smallest significant difference as biologically not meaningful, we concluded that no difference exists.

### Western blotting and densitometry

Lysate was prepared from heart samples which were excised of wound tissue. Isolated heart samples were first washed in PBS and then subjected to lysis with RIPA buffer (50 mM Tris-HCl pH 8.0, 150 mM NaCl, 1 % NP-40, 0.1 % SDS, 0.5 % sodium deoxycholate) using sonication in presence of reducing agents and protease inhibitors DTT, Aprotinin and Leupeptin. Bradford assay for protein estimation, SDS-PAGE and immunoblots were performed according to standard procedures (Bio-Rad system). Nitrocellulose membranes were used for blotting and incubated over night with primary antibodies. Li- COR secondary antibodies were used which were visualized and quantified within a linear range of exposure using a LiCOR ODYSSEY Imager and Image Studio Light software. Bands were quantified using Image Studio Lite Version 5.2. Ratio of p-S6 and p-Akt band intensities to that of the Gapdh loading control of the same blot was calculated. Primary antibodies used were anti-p-S6 (Cell Signaling #2215, 1:1000), anti phospho-serine (473) Akt (Cell signaling #4060S, 1:1000) and anti-Gapdh (Cell signaling #2118, 1:3000). The secondary antibodies were used at 1:10.000 (IRDye conjugates from Li- COR).

### Transmission Electron Microscopy (TEM)

Hearts were fixed in modified Karnovsky’s fixative (2% glutaraldehyde + 2% paraformaldehyde in 50 mM HEPES) for at least overnight at 4°C ^4^ and dissected into small pieces (below 0.5 mm). Samples were washed 2x in 100 mM HEPES and 2x in water and post fixed/contrasted by treatment with osmium tetroxide (OsO_4_), thiocarbohydrazide (TCH), and again OsO_4_ (O-T-O, ^89^. In brief, samples were incubated in 2% aqueous OsO_4_ solution containing 1.5% potassium ferrocyanide and 2 mM CaCl_2_ for 30 min on ice. After washes in water, the samples were incubated in 1% TCH in water (20 min at room temperature), followed by washes in water and a second osmium contrasting step in 2% OsO_4_/water (30 min, on ice). After that, the samples were washed in water, *en bloc* contrasted with 1% uranyl acetate/water for 2 hrs on ice, washed again in water, dehydrated in a graded series of ethanol/water (30%, 50%, 70%, 90%, 96%, 3x 100% ethanol (pure ethanol on molecular sieve)), and infiltrated in epon 812 (epon/ethanol mixtures: 1:3, 1:1, 3:1 for 1.5 hrs each, pure epon overnight, pure epon 5hrs). Finally, the samples were embedded in flat embedding molds and cured at 60°C overnight. Ultrathin sections were prepared with a Leica UC6 ultramicrotome (Leica Microsystems, Vienna, Austria), collected on formvar-coated slot grids, and stained with lead citrate ^90^ and uranyl acetate. Contrasted ultrathin sections were analysed on a Jeol JEM1400 Plus (JEOL, Garching, Germany, camera: Ruby, JEOL) running at 80 kV acceleration voltage.

## Data Availability

The datasets generated during and/or analysed during the current study are available from the corresponding author on reasonable request.

## Acknowledgements

We are thankful to Doris Weber, Brigitte Korte and Martina Raasholm for their excellent fish care and Susanne Kretschmar (EMF, CRTD) for technical assistance. Work in the Weidinger lab was supported by funding from the Deutsche Forschungsgemeinschaft (SFB 1149 [Project Number 251293561], SFB 1279 [Project Number 316249678], WE 4223/8-1 [Project number 433187294], WE 4223/6-1 [Project number 414077062]), by the German ministry of science BMBF (EU ERA-CVD “Cardio-Pro”, grant number 01KL1704), and the Medical faculty of Ulm University (Baustein 3.2 program L.SBN.0168 to M.D.V.) T.K. was supported by the European Regional Development Fund (EFRE, EM and Histology Facility BIOTEC/CRTD), and A.S.P. by the Deutsche Krebshilfe, grant number 70112329.

## Author contributions

A.S.P., Figure 1A, D, E; S.G., Supplementary Figure 6; T.K., Supplementary figure 7; T.B., Supplementary Figure 4E. All other experiments were performed by M.D.V. All authors designed the experiments and analyzed the data. M.D.V. and G.W. wrote the manuscript.

