## Supplementary material for "A tight balance of anabolic mTORC1 signaling and catabolic autophagic activity regulates zebrafish heart regeneration": Supplemtal Figures and Tables

Mohankrishna Dalvoy Vasudevarao[1], Astrid S. Pfister[1], Guoxin Sun[1], Tabea Bieler[1], Alberto Bertozzi[1], Thomas Kurth[2], and Gilbert Weidinger[1,*]

### SUPPLEMENTARY MATERIAL

Supplementary Figure 1 page 2

Supplementary Figure 2 page 3

Supplementary Figure 3 page 4

Supplementary Figure 4 page 5

Supplementary Figure 5 page 7

Supplementary Figure 6 page 8

Supplementary Table 1 page 10

Supplementary Table 2 page 13

### SUPPLEMENTARY FIGURES

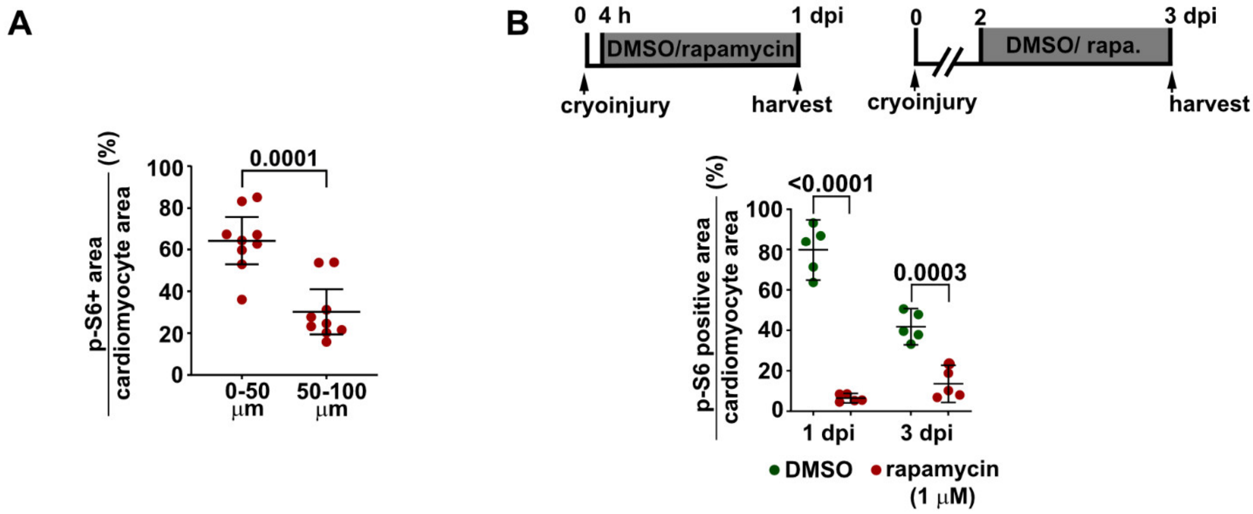


**Supplementary Figure 1: mTOR signaling is activated in border zone cardiomyocytes**.

(A) mTor signaling is more active in Mf20$^+$ cardiomyocytes located at the proximal wound border (0-50 µM) than in the distal wound border (50-100 µm) as revealed by measuring the Mf20$^+$ cardiomyocyte area displaying p-S6 immunoreactivity at 3 dpi. Lines in graph indicate Mean ± C.I. (95%). For further statistical information, see **Supplementary Table 2**.

(B) Treatment with 1 µM rapamycin according to the experimental scheme blocks mTOR activation in cardiomyocytes revealed by p-S6 immunostaining. Scale bar, 100 µm.

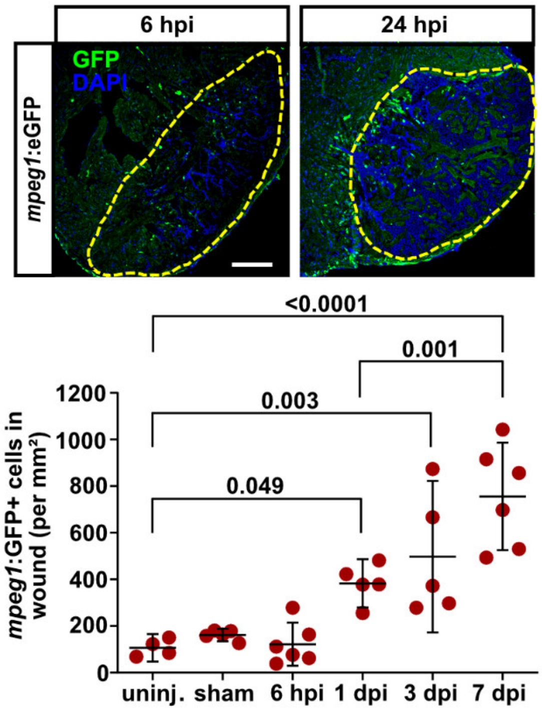


**Supplementary Figure 2: Macrophage recruitment dynamics during heart regeneration.** *mpeg1*:eGFP$^{gl22}$ fish were injured and macrophage recruitment was quantified by counting GFP$^{+}$ cells in the wound area (dashed lines). Significant recruitment to the wound can be detected relative to uninjured hearts starting at 1 dpi. Scale bar, 100 µm. Lines in graph indicate Mean ± C.I. (95%).

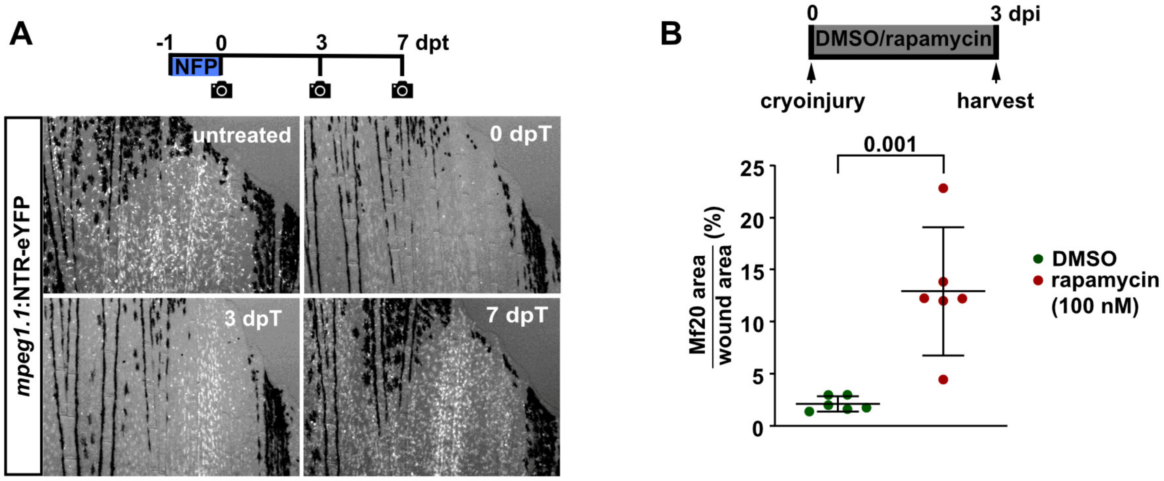


**Supplementary Figure 3**: **Nifurpirinol efficiently ablates macrophages in *mpeg1.1*:NTR-eYFP$^{w202Tg}$ transgenics.**

**(A)** *mpeg1.1*:NTR-eYFP$^{w202Tg}$ transgenics were treated with nifurpirinol (5 μM) for a period of 24 hours and caudal fins were imaged for YFP fluorescence on day 0, 3 and 7 after treatment. A single treatment with nifurpirinol results in drastic reduction of macrophages visible immediately after the end of treatment (0 dpT). Subsequently, by day 3 macrophages substantially repopulate the fins and by day 7 appear to have reached normal levels.

**(B)** Rapamycin treatment (100 nM) causes accumulation of Mf20$^+$ debris by 3 dpi in cryoinjured zebrafish hearts. Lines in graph indicate Mean ± C.I. (95%).

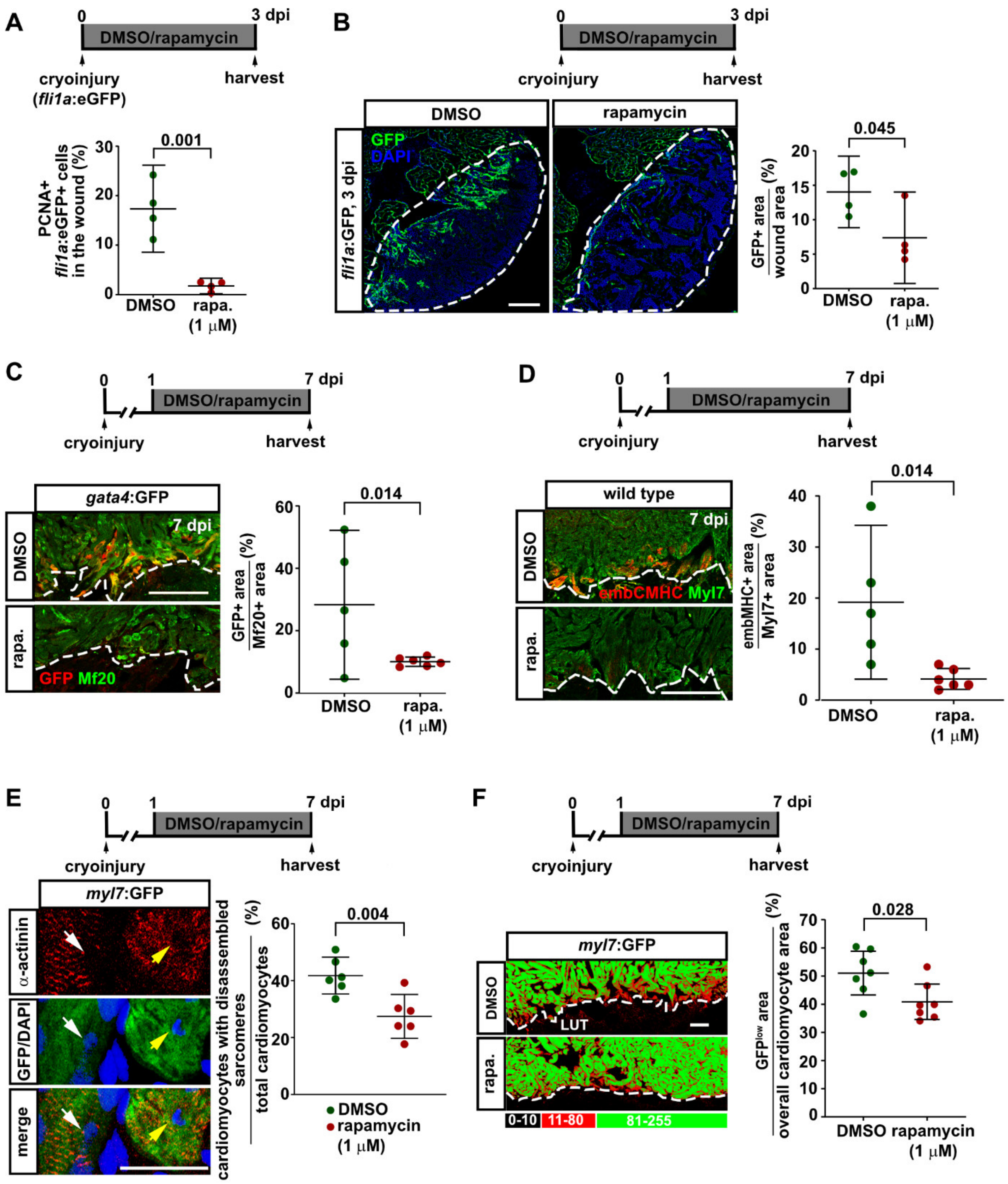


**Supplementary Figure 4: mTOR signaling is required for wound re-vascularization and cardiomyocyte dedifferentiation.**

(**A**) Treatment with rapamycin (1 μM) reduces the number of PCNA+ endothelial (*flia*:eGFP+) cells in the wound at 3 dpi. Lines in graph indicate Mean ± C.I. (95%).

(**B**) A plexus of *fli1a*:eGFP$^{y1Tg+}$ endothelial cells has formed proximal to the wound border in the wound of control hearts, but less so in rapamycin treated (1 μM) hearts. The area occupied by *flia*:eGFP+ cells relative to the wound area is plotted. Scale bar, 100 μm.

(**C**) Rapamycin treatment (1 μM) blocks upregulation of *gata4* regulatory sequences in $Mf20^+$ cardiomyocytes of *14.8gata4*:$GFP^{ae1tg}$ transgenic fish at 7 dpi. Size of the $GFP^+$ area relative to the wound border $Mf20^+$area is plotted. Scale bar, 100 μm.

(**D**) Immunostaining against an embryonic cardiac MHC isoform (embCMHC) detected by the N2.261 antibody and Myl7 reveals that treatment with rapamycin (1 μM) for 6 days suppresses re-expression of embCMHC in the wound border myocardium (0-50 μm) in injured hearts at 7 dpi. The $embCMHC^+$ area relative to the $Myl7^+$ area in the wound border myocardium is plotted. Scale bar, 100 μm.

(**E**) Immunostaining against α-actinin reveals normal sarcomeres (white arrow) or disorganized sarcomeres (yellow arrow) surrounding nuclei of *myl7*:$GFP^{twuf1tg\ +}$ cardiomyocytes in border zone myocardium (0-50 μm) at 7 dpi. Treatment with 1 μM rapamycin reduces the number of cardiomyocytes displaying disassembled sarcomeres, plotted relative to the total number of cardiomyocytes within 50 μm of the wound border. Scale bar, 50 μm.

(**F**) LUT (lookup table) images displaying *myl7*:$GFP^{twu34Tg}$ fluorescent intensity below ~30% of maximum intensity in red and the rest in green. Rapamycin treatment (1 μM) from 1 until 7 dpi reduces the area occupied by low GFP expressing cells (red) relative to the overall myocardial area at the wound border (0-50 μm). Scale bar, 50 μm.

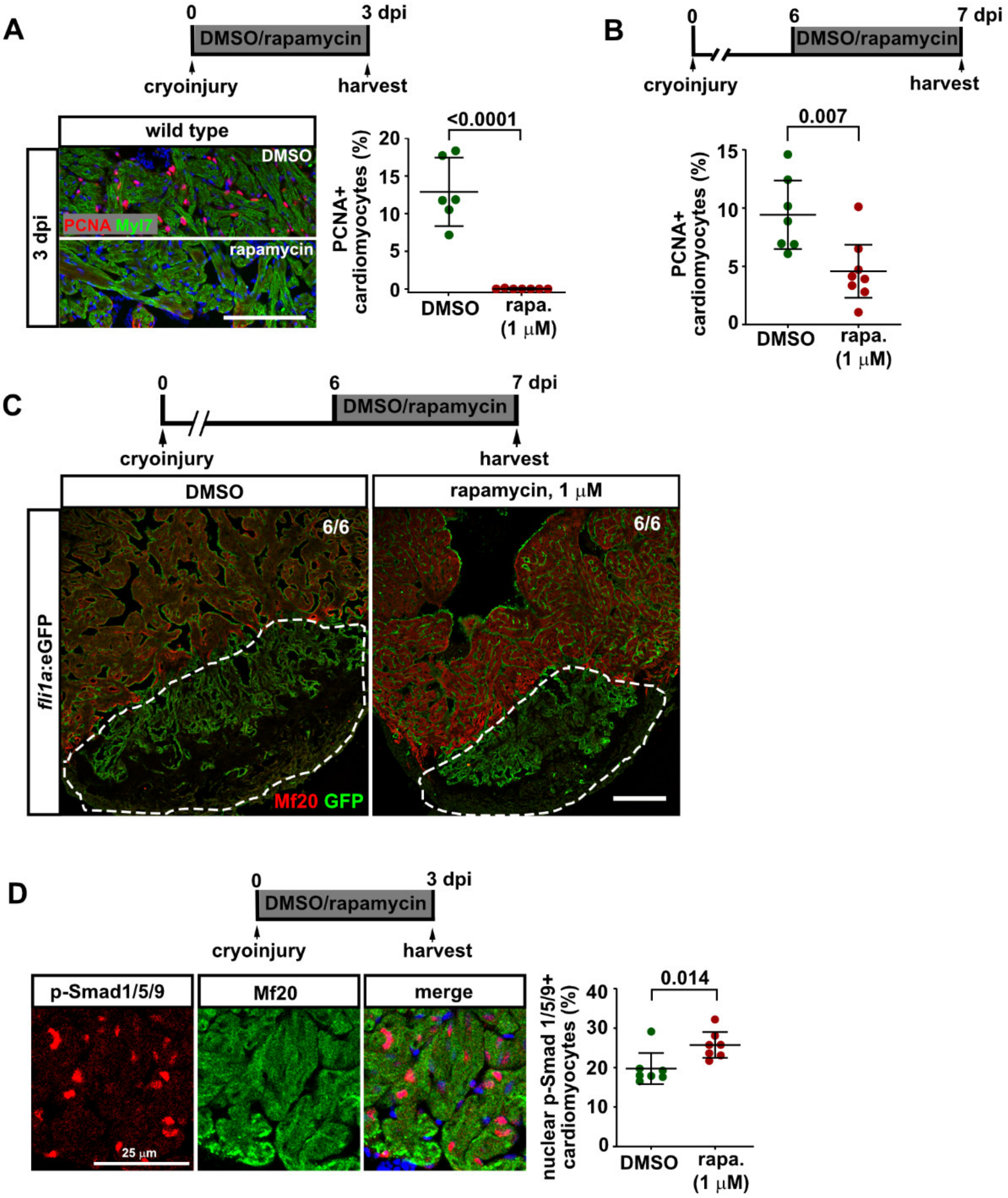


**Supplementary Figure 5: mTOR signaling is required for cardiomyocyte proliferation but not for BMP signaling activation.**

(**A**) Rapamycin treatment (1 μM) for 3 days severely reduces the number of PCNA$^+$ Myl7$^+$ cardiomyocytes at the wound border (0-100 μm) at 3 dpi. Scale bar, 100 μm. Lines in graph indicate Mean ± C.I. (95%).

(**B**) 24 hours of rapamycin treatment (1 μM) reduces cardiomyocyte cell cycle activity at 7 dpi.

(**C**) GFP immunostaining in *fli1a*:eGFP$^{y1Tg}$ hearts at 7 dpi treated from 6-7 dpi with either DMSO or rapamycin reveals similar extent of vascular plexus formation (GFP$^+$ area within the wound indicated by dashed lines). Images are representative for 6/6 hearts for each group. Scale bar, 100 μm.

(**D**) SMAD1/5/9 accumulates in the nucleus of Mf20$^+$ cardiomyocytes at the wound border at 3 dpi. Treatment with rapamycin (1 μM) for 3 days does not reduce the number of p-SMAD1/5/9$^+$ wound border (0-100 μm) cardiomyocytes, but rather slightly increases it. Scale bar, 25 μm.

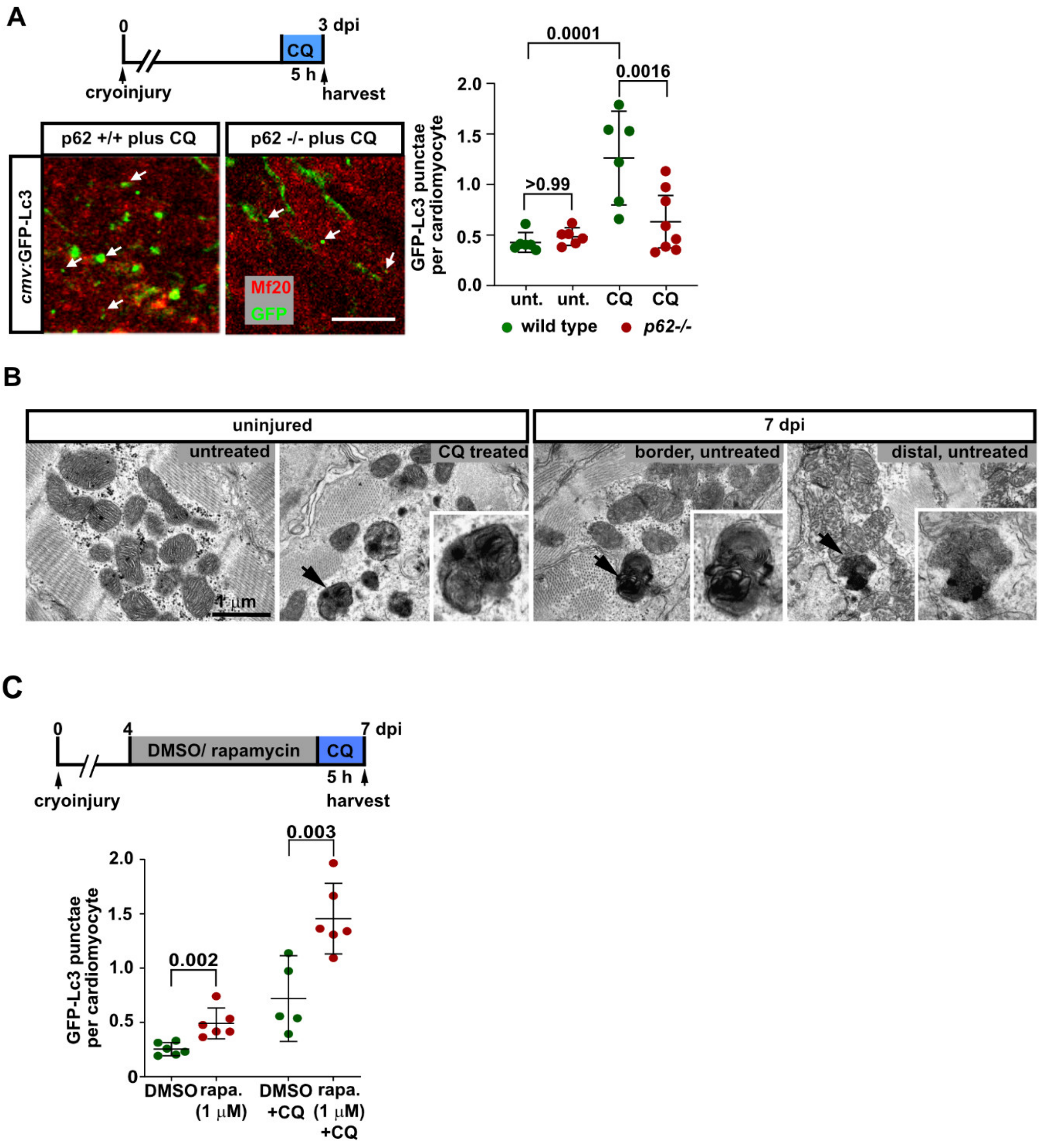


**Supplementary Figure 6: Autophagy is upregulated during heart regeneration while mTOR signaling limits it in the border zone cardiomyocytes.**

(**A**) At 3 dpi, GFP-Lc3$^{+}$ punctae were quantified in Mf20$^{+}$ cardiomyocytes located at the wound border (0-100 µm) in *cmv*:GFP-Lc3 reporter fish wild-type for *p62* or homozygous for the *p62*$^{ibl52}$ loss of function allele. Treatment with 0.8 mM of the lysosomal inhibitor chloroquine (CQ) for 5 hours increases the number of punctae in wild type but not in mutant hearts. Lines in graph indicate Mean ± C.I. (95%). Scale bar, 10 µm.

(**B**) Autophagosomes characterized by bilamellar appearance (arrows) can be detected by transmission electron microscopy in the myocardium of injured hearts at 7 dpi both in the wound border and distal areas, but in uninjured hearts only after treatment with chloroquine. Scale bar, 1 µm.

(**C**) The number of GFP-Lc3$^+$ punctae detected per cell in wound border (0-100 μm) Mf20$^+$ cardiomyocytes at 7 dpi is increased upon treatment with 1 μM rapamycin for 3 days. Treatment with 0.8 mM chloroquine for 5 hours is sufficient to increase the number further in both control and rapamycin treated hearts.

### SUPPLEMENTARY TABLES

#### Supplementary Table 1.
#### Additional statistical information about data presented in Main Figures 1-7.

| Figure | Subfigure | Groups | **sample no.** (hearts, unless specified otherwise) | Statistical test | ANOVA p value |
|---|---|---|---|---|---|
| **Figure 1** | 1(A) | Uninjured | 4 lysates (each pooled from 10 ventricles) | paired 1-tailed t- test | - |
| | | 3 dpi | 4 lysates (each pooled from 10 ventricles) | | |
| | 1(B) | Uninjured | 9 | one way ANOVA Bonferroni´s multiple comparisons test | <0.0001 |
| | | 1 dpi | 10 | | |
| | | 3 dpi | 10 | | |
| | | 7 dpi | 10 | | |
| | | 14 dpi | 10 | | |
| | 1(C) | uninjured | 5 | unpaired 2-tailed t test | |
| | | 6 hpi | 5 | | |
| | 1(D) | 1 dpi DMSO | 6 | unpaired 2-tailed t test | - |
| | | 1 dpi rapamycin | 6 | | |
| | 1(E) | uninjured | 4 lysates (each pooled from 10 ventricles) | paired 1-tailed t test | - |
| | | 1 dpi DMSO | 4 lysates (each pooled from 10 ventricles) | | |
| | | 1 dpi rapamycin | 4 lysates (each pooled from 10 ventricles) | | |
| | | 1 dpi torin1 | 4 lysates (each pooled from 10 ventricles) | | |
| | 1(F) | uninjured | 5 lysates (each pooled from 7 ventricles) | paired 1-tailed t test | - |
| | | 3 dpi | 5 lysates (each pooled from 7 ventricles) | | |
| | 1(G) | uninjured | 4 | one way ANOVA Bonferroni´s multiple comparisons test | <0.0001 |
| | | 3 dpi, wound, | 4 | | |
| | | 3 dpi, border, | 4 | | |

| Figure | Subfigure | Groups | **sample no.** (hearts) | Statistical test | ANOVA p value |
|---|---|---|---|---|---|
| **Figure 2** | 2(A) | uninjured | 6 | one way ANOVA Bonferroni´s multiple comparisons test | 0.0006 |
| | | 1 dpi border | 11 | | |
| | | 1 dpi distal | 11 | | |
| | | 3 dpi border | 12 | | |
| | | 3 dpi distal | 12 | | |
| | 2(B) | ethanol | 10 | unpaired 2-tailed t test | - |
| | | dexamethasone | 10 | | |

| Figure | Subfigure | Groups | sample no. (hearts) | Statistical test | ANOVA p value |
|---|---|---|---|---|---|
| Figure 3 | 3(B) | uninjured | 5 | one way ANOVA Bonferroni´s multiple comparisons test | <0.0001 |
| | | sham | 4 | | |
| | | 6 hpi | 6 | | |
| | | 1 dpi | 6 | | |
| | | 3 dpi | 5 | | |
| | | 7 dpi | 6 | | |
| | | rapamycin | 4 | | |
| | 3(C) | DMSO | 12 | one way ANOVA Bonferroni´s multiple comparisons test | <0.0001 |
| | | torin1 | 11 | | |
| | | rapamycin | 12 | | |
| | 3(D) | uninjured | 6 | one way ANOVA Bonferroni´s multiple comparisons test | <0.0001 |
| | | sham | 6 | | |
| | | 1 dpi | 6 | | |
| | 3(E) | uninjured sham | 5 | one way ANOVA Bonferroni multiple comparisons test | 0.336 |
| | | sham | 6 | | |
| | | 1 dpi | 6 | | |
| | 3(F) | DMSO | 6 | unpaired 2-tailed t test | - |
| | | rapamycin | 5 | | |

| Figure | Subfigure | Groups | sample no. (hearts) | Statistical test | ANOVA p value |
|---|---|---|---|---|---|
| Figure 4 | 4(B) | PBS liposomes | 6 | unpaired 2-tailed t test | - |
| | | clodrosomes | 6 | | |
| | 4(C) | PBS liposomes | 6 | unpaired 2-tailed t test | - |
| | | clodrosomes | 6 | | |
| | 4(D) | Wild type +NFP | 7 | unpaired 2-tailed t test | - |
| | | *mpeg1.1*:NTR-YFP | 7 | | |
| | 4(E) | 1 dpi DMSO | 5 | one way ANOVA Bonferroni multiple comparisons test | <0.0001 |
| | | 1 dpi rapamycin | 5 | | |
| | | 3 dpi DMSO | 5 | | |
| | | 3 dpi rapamycin | 5 | | |
| | 4(F) | 3 dpi rapamycin | 12 | one way ANOVA Bonferroni multiple comparisons test | 0.0001 |
| | | 5 dpi rapamycin | 11 | | |
| | | 5 dpi rapamycin+NFP | 12 | | |
| | 4(G) | 1 dpi | 14 | unpaired 2-tailed t test | - |
| | | 3 dpi | 15 | | |

| Figure | Subfigure | Groups | sample no. (hearts) | Statistical test | ANOVA p value |
|---|---|---|---|---|---|
| Figure 5 | 5(A) | DMSO | 7 | unpaired 2-tailed t test | - |
| | | rapamycin | 7 | | |
| | 5(B) | DMSO | 6 | unpaired 2-tailed t test | - |
| | | rapamycin | 6 | | |
| | 5(C) | DMSO | 9 | unpaired 2-tailed t test | - |
| | | Rapamycin | 10 | | |
| | 5(D) | 3 dpi, DMSO | 7 | one way ANOVA Bonferroni multiple comparisons test | 0.0067 |
| | | 18 dpi, DMSO | 9 | | |
| | | 18 dpi, rapamycin | 8 | | |
| | 5(E) | 7 dpi, DMSO | 6 | one way ANOVA Bonferroni multiple comparisons test | <0.0001 |
| | | 21 dpi, DMSO | 7 | | |
| | | 21 dpi, rapamycin | 8 | | |

| Figure | Subfigure | Groups | sample no. (hearts) | Statistical test | ANOVA p value |
|---|---|---|---|---|---|
| **Figure 6** | 6(A) | uninjured | 5 | one way ANOVA Bonferroni multiple comparisons test | <0.0001 |
| | | 1 dpi | 5 | | |
| | | 3 dpi | 5 | | |
| | | 7 dpi | 6 | | |
| | | 14 dpi | 6 | | |
| | | 21 dpi | 6 | | |
| | 6(B) | uninjured | 5 | one way ANOVA Bonferroni multiple comparisons test | <0.0001 |
| | | uninjured chloroquine | 7 | | |
| | | 3 dpi, border, untreated | 6 | | |
| | | 3 dpi, border chloroquine | 7 | | |
| | | 3 dpi, distal, untreated | 6 | | |
| | | 3 dpi, distal, chloroquine | 7 | | |
| | 6(C) | 1 dpi | 5 | unpaired 2-tailed t test | - |
| | | 3 dpi | 5 | | |
| | 6(D) | DMSO | 6 | unpaired 2-tailed t test | - |
| | | rapamycin | 6 | | |

| Figure | Subfigure | Groups | sample no. (hearts) | Statistical test | ANOVA p value |
|---|---|---|---|---|---|
| **Figure 7** | 7(A) | DMSO border | 6 | one way ANOVA Bonferroni multiple comparisons test | <0.0001 |
| | | MEK1-I border | 6 | | |
| | | DMSO distal | 6 | | |
| | | MEK1-I distal | 6 | | |
| | 7(B) | DMSO border | 12 | one way ANOVA Bonferroni multiple comparisons test | <0.0001 |
| | | JNK-i border | 13 | | |
| | | DMSO distal | 11 | | |
| | | JNK-I distal | 13 | | |
| | 7(C) | DMSO | 6 | one way ANOVA Bonferroni multiple comparisons test | <0.0001 |
| | | MEK1-i | 6 | | |
| | | DMSO plus chloroquine | 6 | | |
| | | MEK1-I plus chloroquine | 6 | | |
| | 7(D) | DMSO | 7 | one way ANOVA Bonferroni multiple comparisons test | <0.0001 |
| | | JNK-i | 7 | | |
| | | DMSO plus chloroquine | 6 | | |
| | | JNK-I plus chloroquine | 6 | | |
| | 7(E) | 3 dpi DMSO | 10 | Unpaired 2-tailed t test | - |
| | | 3 dpi MEK1-i | 10 | | |

**Supplementary Table 2.**
**Additional statistical information about data presented in Supplementary Figures.**

| Figure | Subfigure | Group | sample no. (hearts) | statistical test | ANOVA p-value |
|---|---|---|---|---|---|
| S1 | S1(A) | 0-50 micrometers | 9 | Unpaired 2-tailed t test | - |
| | | 50-100 micrometers | 9 | | |
| | S1(B) | 1 dpi, DMSO | 5 | one way ANOVA Bonferroni multiple comparisons test | <0.0001 |
| | | 1 dpi, rapamycin | 5 | | |
| | | 3 dpi, DMSO | 5 | | |
| | | 3 dpi, rapamycin | 5 | | |
| S2 | - | uninjured | 4 | Unpaired 2-tailed t test | - |
| | | Sham | 4 | | |
| | | 6 hpi | 6 | | |
| | | 1 dpi | 5 | | |
| | | 3 dpi | 5 | | |
| | | 7 dpi | 6 | | |
| S3 | S3(B) | DMSO | 6 | Unpaired 2-tailed t test | - |
| | | rapamycin | 6 | | |
| S4 | S4(A) | DMSO | 4 | Unpaired 2-tailed t test | - |
| | | rapamycin | 4 | | |
| | S4(B) | DMSO | 4 | Unpaired 2-tailed t test | <0.0001 |
| | | rapamycin | 4 | | |
| | S4(C) | DMSO | 5 | Unpaired 2-tailed t test | |
| | | rapamycin | 6 | | |
| | S4(D) | DMSO | 5 | Unpaired 2-tailed t test | |
| | | rapamycin | 6 | | |
| | S4(E) | DMSO | 6 | Unpaired 2-tailed t test | |
| | | rapamycin | 6 | | |
| | S4(F) | DMSO | 7 | Unpaired 2-tailed t test | |
| | | rapamycin | 7 | | |
| S5 | S5(A) | DMSO | 6 | unpaired 2-tailed t test | - |
| | | rapamycin | 7 | | |
| | S5(B) | DMSO | 7 | unpaired 2-tailed t test | - |
| | | rapamycin | 8 | | |
| | S5(D) | DMSO | 7 | unpaired 2-tailed t test | - |
| | | rapamycin | 7 | | |
| S6 | S6(A) | wild type untreated | 6 | one way ANOVA Bonferroni multiple comparisons test | 0.0001 |
| | | p62 homozygous, untreated | 6 | | |
| | | wild type, chloroquine | 6 | | |
| | | p62 homozygous, chloroquine | 8 | | |
| | S6(C) | DMSO | 6 | one way ANOVA Bonferroni multiple comparisons test | <0.0001 |
| | | rapamycin | 6 | | |
| | | DMSO + chloroquine | 5 | | |
| | | Rapamycin +chloroquine | 6 | | |
